# Rho1 and Rgf3 regulate the expansion of the nuclear envelope during fission yeast mitosis/cytokinesis

**DOI:** 10.64898/2026.06.22.733743

**Authors:** Rubén Celador, Patricia García, Virginia Tajadura, Tomás Edreira, Mireia Casasampere, James B. Moseley, Yolanda Sánchez

## Abstract

The nuclear envelope (NE) surrounds the genetic material and is continuous with the endoplasmic reticulum (ER). In yeast and other organisms undergoing closed mitosis, nuclear envelope expansion (NME) is strictly required to accommodate spindle elongation and ensure proper chromosome segregation within a single nuclear compartment. Failure to expand the NE during mitosis leads to chromosome missegregation. Here, we show that deletion of the unstructured N-terminal domain of Rgf3, a Rho1-specific guanine nucleotide exchange factor (GEF), causes early mitotic defects that produce the characteristic “cut” phenotype of untimely cell division. The *rgf3*ΔN2 mutant displays spindle buckling, a hallmark of anaphase nuclei unable to properly expand the NE. From yeast to mammals, phosphatidic acid (PA)—a key precursor in phospholipid biosynthesis—is metabolized via two competing pathways, the cytidine diphosphate–diacylglycerol (CDP–DAG) and the Kennedy pathways, both contributing to lipid membrane homeostasis. We provide evidence that impaired Rho1 activation in *rgf3*ΔN2 selectively disrupts phospholipid synthesis through the CDP-choline branch of the Kennedy pathway. Thus, Rho1 promotes mitotic progression by modulating phospholipid biosynthesis to enable efficient NME during anaphase.

**Highlights:** The N-terminus of Rgf3 is required for proper nuclear envelope expansion (NME) during anaphase.

The structurally flexible N-terminal domain of Rgf3 is essential for localized Rho1 activation.

Active Rho1 drives mitotic membrane growth by modulating phospholipid synthesis through the Kennedy pathway.

## Introduction

Mitosis in eukaryotes requires remodeling of the nuclear envelope (NE) to enable chromosome segregation by the microtubule-based spindle ^1–5^. In open mitosis, which is characteristic of metazoans and plants cells, the NE retracts into the endoplasmic reticulum (ER) during NE breakdown, allowing spindle assembly to occur directly in the cytoplasm ^6,7^. By contrast, cells undergoing closed mitosis, such as budding and fission yeast, must import spindle components into the nucleus, assemble spindle pole bodies (SPBs), and expand the NE to accommodate spindle elongation ^8–10^.

In mammalian cells, NE reassembly after chromosome segregation depends on the incorporation of newly synthesized phospholipids into membranes that reassociate with and enclose the segregated chromatin ^11^. Similarly, mitotic NE elongation in yeasts requires an increase in glycerophospholipid (GPL) biosynthesis to sustain membrane growth ^4,12,13^. GPL biosynthesis needed for membrane expansion is regulated at the metabolic branch point defined by the reversible conversion of phosphatidic acid (PA) to diacylglycerol (DAG). PA can be converted into the GPL precursor cytidine diphosphate–diacylglycerol (CDP-DAG) or dephosphorylated to DAG by lipin phosphatases. DAG, in turn, serves as a substrate for triacylglycerol (TAG) synthesis and to produce the major membrane lipids phosphatidylcholine (PC) and phosphatidylethanolamine (PE) via the Kennedy pathway ^14,15^.

In budding and fission yeast, expansion of the NE is driven by the CDK1-dependent inactivation and delocalization of lipin phosphatases (Pah1 in *S. cerevisiae* and Ned1 in *S. pombe*). When lipin is inactivated upon mitotic entry, the block in PA-to-DAG conversion increases GPL biosynthesis, promoting nuclear/ER membrane expansion ^13,16^. Lipin activity is an important point of regulation. Lipin is normally stimulated by the Nem1/Spo7 phosphatase complex ^13,16–18^, which is itself inhibited by SUMOylation, promoting Nem1/Spo7 dissociation from membranes ^19^, and by ER resident proteins, among others ^20^. Conversion of DAG to PA by the diacylglycerol kinase Dgk1 provides an additional layer of control, though with species-specific consequences ^20–22^. In *S. cerevisiae*, loss of Dgk1 causes only moderate defects in nuclear morphology, whereas in *S. pombe* more than 40% of *dgk1*Δ cells fail to expand the nuclear membrane during mitosis ^23^. Moreover, NE expansion in *S. pombe* also requires that DAG be funneled into GPL synthesis through the Kennedy pathway ^23^.

The Rho family of GTPases are membrane-associated molecular switches that regulate cytoskeletal dynamics. In animal cells, Rho GTPases (RhoA, Rac1, and Cdc42) regulate changes in cell shape by controlling actin nucleation complexes in response to a variety of stimuli ^24^. This remodeling can result in an increase or decrease in membrane surface area and tension, driven by the fusion (exocytosis) or internalization (endocytosis) of membranous compartments, respectively ^25^. Similar to animal cells, budding yeast Rho1 is a critical regulator of extracellular matrix remodeling, as it controls the activities of Fsk1 (β-glucan synthase) and Chs3 (chitin synthase 3), and regulates the localization of the exocyst complex ^26,27^. In fission yeast, Rho1-dependent actin localization is required for the correct positioning of growth poles ^28^. Rho1 activates cell wall synthesis and contributes to the preservation of membrane integrity ^29,30^. However, the function of Rho1 at the early stages of cytokinesis is largely unknown ^31,32^. In dividing cells, Rho1 localizes to perinuclear membranes and the cleavage furrow, and is activated by three GEFs: Rgf1, Rgf2, and Rgf3 ^31^. Rgf1 signal is excluded from the nucleus during interphase ^33^, but, in anaphase, assembles into a ring that migrates centripetally with the membrane that seeds the growing septum ^30,34,35^. Rgf1 also functions in a checkpoint-like pathway that delays actomyosin ring (CAR) constriction initiation under cell wall stress ^36,37^. Rgf2 acts specifically during sporulation, with no established role in division ^37^. By contrast, Rgf3 forms a ring early in anaphase (when SPBs are ∼3µm apart) that contracts with the CAR ^34,35,38^, positioned between the membrane-associated scaffolds Mid1, Cdc15, and Imp2 externally, and F-actin with associated motors internally ^39,40^. Rgf3 stimulates Rho1-mediated activation of β-glucan synthase activity ^38^ and interacts with the arrestin Art1, which most likely connects Rgf3 to receptors of the endocytic machinery ^41^. Surprisingly, we found that deletion of the N-terminus of Rgf3 does not produce cell wall defects but instead leads to nuclear envelope shrinkage and spindle bending that is exacerbated upon perturbation of nuclear membrane synthesis. The unstructured N-terminus of Rgf3 is crucial for Rho1 activation promoting membrane expansion. We propose that Rgf3 and Rho1 could be key players in the mechanism regulating the explosion of phospholipid synthesis necessary to achieve NM outgrowth ahead of cell separation.

## Results

### The N-terminus of Rgf3 is required for proper nuclear division

To explore the role of Rgf3 in cell division, we generated a series of N- and C-terminal deletion mutants, excluding the DH and PH domains known to be essential for catalysis (Figure S1A). Cells harboring N-terminal truncations were morphologically indistinguishable from wild-type (WT) cells, and the mutant proteins exhibited normal sensitivity to cell wall-related stresses (Figure S1A and S1B). We chose the *rgf3-*ΔN2 mutant —which lacks the entire N-terminus (amino acids [aa]11-295)— for further characterization. We next examined contractile actomyosin ring (CAR) dynamics via time-lapse fluorescence microscopy in cells expressing Imp2-GFP (CAR marker) and Sfi1-GFP (SPB marker) (Figure 1A). Neither WT nor *rgf3*ΔN2 early anaphase cells displayed structural abnormalities in the actomyosin rings. Moreover, the total duration of cytokinesis —measured from the onset of SPB separation to the completion of CAR constriction— was statistically comparable, lasting on average 33.88 ± 2.39 min in WT cells compared to 34.2 ± 2.82 minutes (min) in *rgf3-*ΔN2 mutants (Figure 1B). Nonetheless, a significant fraction of *rgf3-*ΔN2 cells (∼15%) displayed anomalous SPB segregation (Figure 1C). In these cases, the SPBs initially separated normally during early anaphase; however, instead of continuing their elongation toward the cell poles, they reversed direction, moved toward one another, and frequently ended up together within a single daughter cell (Figure 1A). To monitor nuclear dynamics during these faulty divisions, we visualized the nuclear mass using Hht1-RFP (histone H3). We observed two distinct types of abnormal segregation in *rgf3*ΔN2 mutants: anucleated cells resulting from septation in cells with displaced nuclei (Figure 1D, cell a), and uninucleated cells containing nuclei of asymmetrical sizes, due to biased nuclear positioning prior to cytokinesis (Figure 1D, cell b). Both phenotypes occurred at similar frequencies. Indeed, septation itself was not impeded in these mutants, as cell wall biosynthesis—tracked by Blankophor (BP) staining—remained unaltered (Figure 1E).

**Figure 1.**
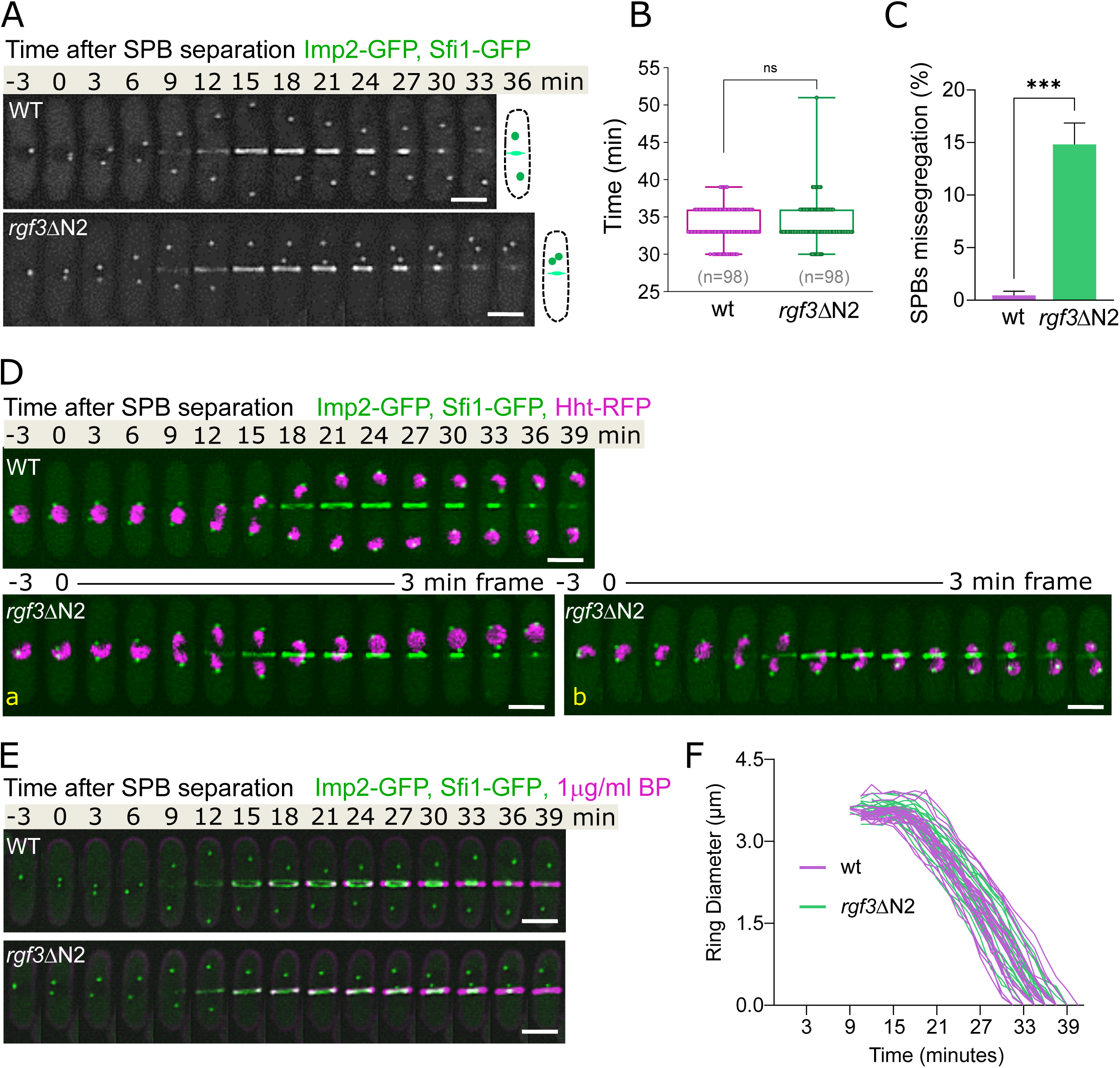
The N-terminus of Rgf3 is required for proper chromosomal segregation. **(A)** Representative time-lapse fluorescence micrographs of wild-type (WT) and *rgf3*ΔN2 cells expressing Imp2-GFP (CAR marker) and Sfi1-GFP (SPB marker). Diagrams on the right illustrate the representative terminal phenotype for each strain. **(B)** Quantification of cytokinesis duration (from SPB separation to the end of CAR contraction) in cells from (A) (n = 98 cells per strain). Boxes represent the interquartile range (IQR), whiskers depict Tukey’s range, and the horizontal line inside the box indicates the median value (n.s., non-significant, calculated by an unpaired Student’s t-test). **(C)** Percentage of SPB missegregation in WT and *rgf3*ΔN2 cells. Data represent the mean ± SD from three independent experiments (n > 20 cells analyzed per strain; ***, p < 0.0001, calculated by an unpaired Student’s t-test). **(D)** Time-lapse series of WT and *rgf3*ΔN2 cells expressing Imp2-GFP, Sfi1-GFP, and Hht1-RFP (histone H3; nuclear mass marker). For the *rgf3*ΔN2 mutant, two representative examples of common missegregation phenotypes are shown (cell “a”, left; cell “b”, right). All cells were cultured at 28°C; images are maximum-intensity projections of Z-stacks. Scale bars, 5 µm. **(E)** Representative time-lapse micrographs of wild-type (WT) and *rgf3*ΔN2 cells expressing Imp2-GFP (CAR marker) and Sfi1-GFP (SPB marker). Cells were cultured at 28°C and imaged in the presence of Blankophor (1µg/ml) added immediately prior to imaging for cell wall/septum visualization. Quantification of cytokinesis kinetics from these cells is presented in Figure S2A. Scale bars, 5 µm. **(F)** Time course of CAR constriction in WT and *rgf3*ΔN2 cells expressing Imp2-GFP and Sfi1-GFP. The graph displays the ring diameter over time; with time 0 min defined as SPB separation; curves represent individual cells (n = 25) per strain.

We hypothesized that a premature initiation of septation in *rgf3*ΔN2 cells might drive these segregation defects. However, we detected no differences in key cytokinetic parameters. Initial Imp2-GFP rings in WT and *rgf3*ΔN2 cells undergoing missegregation exhibited similar diameters (WT: 3.62 ± 0.16; *rgf3*ΔN2: 3.65 ± 0.16 µm) and constricted at identical rates ∼0.18 µm/min (Figures 1F, S1C and S1D). Furthermore, the mean time to the onset of ring constriction was equivalent between both strains (Figures 1F and S1E). We also evaluated whether the Septation Initiation Network (SIN) pathway was prematurely activated in *rgf3*ΔN2 cells by monitoring the asymmetric recruitment of Cdc7-GFP to the SPBs ^42–44^. Cdc7-GFP asymmetry was established normally and peaked concurrently in both strains (Figure S1F). As expected, in *rgf3*ΔN2 cells exhibiting a “cut” phenotype—where the septum constricts but nuclear separation fails—the asymmetric SIN signaling persisted aberrantly (Figure S1F). In conclusion, in a significant percentage of *rgf3-*ΔN2 cells entering mitosis, the chromosomes were intersected by the septum or displaced off-center during cytokinesis. This failure, however, is not a consequence of premature initiation of septation or untimely SIN signaling.

### The absence of the N-terminus of Rgf3 promotes spindle buckling during anaphase

The “cut”-like phenotype is frequently associated with mutations in genes involved in anaphase promotion and chromosome segregation ^45–48^. Therefore, we examined the metaphase-to-anaphase transition in *rgf3*ΔN2 cells exhibiting SPB missegregation. Utilizing live-cell imaging, we tracked the distance between SPBs over time to determine spindle length kinetics. Spindle elongation in wild-type (WT) cells is classically described as a three-phase process: Phase I corresponds to spindle assembly during pro-metaphase, characterized by a low elongation velocity; Phase II represents metaphase and anaphase A, marked by a relatively constant spindle length; and Phase III corresponds to anaphase B, which is driven by rapid spindle elongation (Figure 2A and 2B). No significant differences in spindle length were observed between WT and *rgf3*ΔN2 cells during Phases I and II. Indeed, both strains reached a comparable steady-state metaphase spindle length of 2.39 ± 0.47 µm and 2.45 ± 0.25 µm, respectively, over a period of approximately 9 minutes (Figure 2A and S2B). However, *rgf3*ΔN2 cells exhibited a significant and persistent reduction in spindle length during Phase III (Figures 2B and 2C). Mutant cells undergoing missegregation assembled spindles that reached a maximum length of only 5.43 ± 0.31 µm, compared to an average maximum length of 10.60 ± 0.44 µm in WT control cells (Figure 2B). These findings suggests that *rgf3*ΔN2 cells fail to successfully complete Phase III spindle elongation.

**Figure 2.**
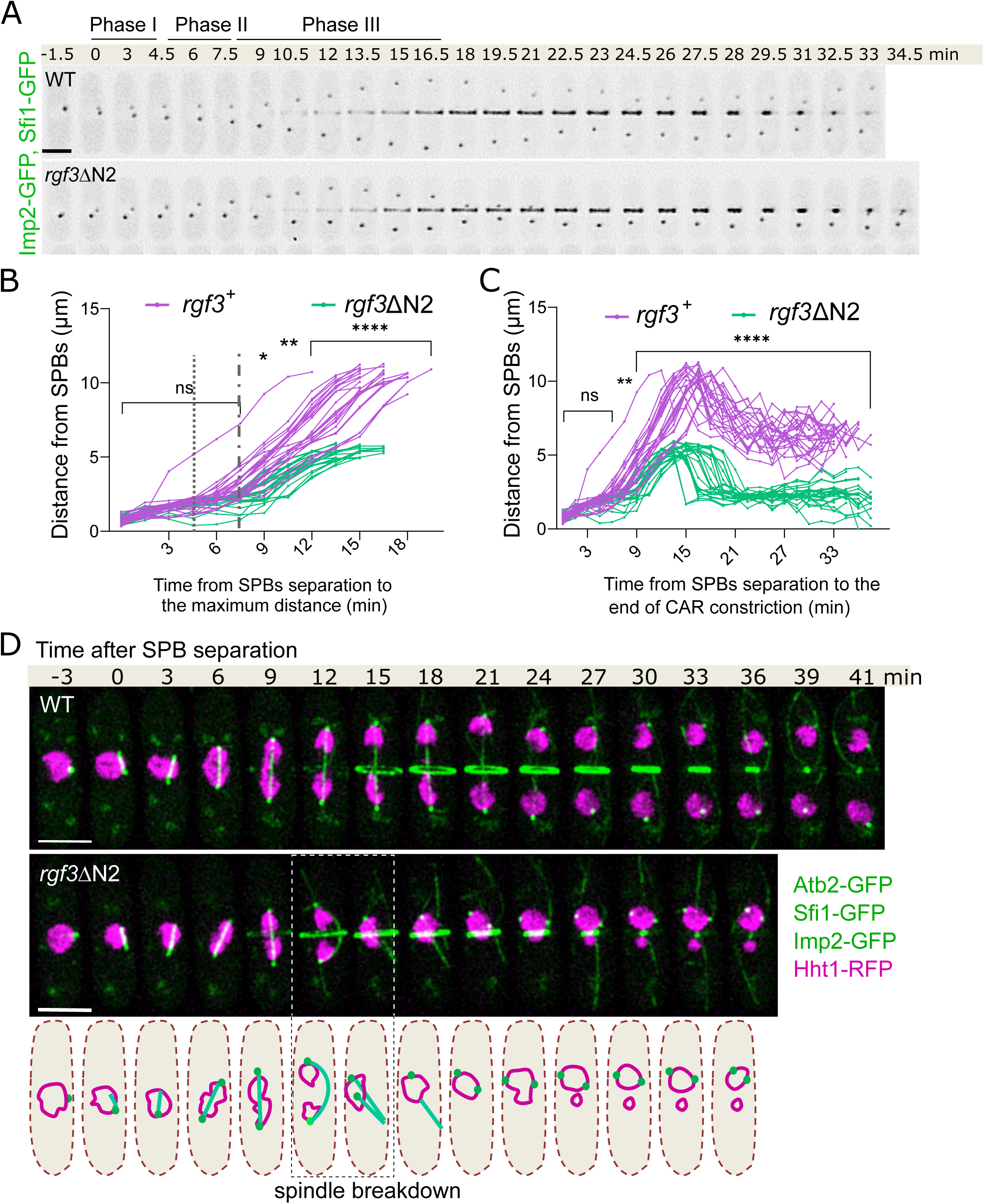
The rgf3-ΔN2 mutation promotes spindle buckling in anaphase. **(A)** Representative time-lapse micrographs of wild-type and *rgf3*ΔN2 cells expressing Imp2-GFP (CAR marker) and Sfi1-GFP (SPB marker). Images are inverted grayscale maximum-intensity projections of seven Z-sections captured at the indicated time points (min); time 0 min corresponds to SPB separation. The developmental stages of the spindle are categorized into Phase I, Phase II, and Phase III (based on spindle elongation kinetics). Scale bar, 5 µm. **(B)** Comparative plot of spindle length dynamics in *rgf3*^+^ (purple curves, n=25) and *rgf3*ΔN2 cells (green curves, n=25), measured as the distance between SPBs over time. The dotted vertical line marks the phase I–II transition, whereas the dashed vertical line indicates the Phase II–III transition. Asterisks denote statistical significance between the two genotypes at specific time points, calculated by ANOVA followed by Šidák’s multiple comparisons test, calculated using ANOVA followed by the Šidák multiple comparisons test (n.s. p>0.05; * p<0.05; **, p<0.01; ****, p<0.0001). **(C)** Distance between SPBs over time from SPB separation to the end of CAR constriction in (purple curves, n=25) and cells (green curves, n=25). The distance between the SPBs in wild type and *rgf3*-ΔN2 cells from (B) was plotted from the onset of SPB separation until the end of CAR constriction. Asterisks denote statistical significance between genotypes, calculated by ANOVA followed by Šidák’s multiple comparisons test (n.s. p>0.05; **, p<0.01; ****, p<0.0001). **(D)** Representative time-lapse fluorescence micrographs of wild-type and *rgf3*ΔN2 cells expressing Atb2-GFP (spindle marker), Imp2-GFP (CAR marker), Sfi1-GFP (SPB marker) and Hht1-RFP (nuclear mass marker). The dashed box highlights the spindle collapse observed in *rgf3*ΔN2 cells. The schematic diagram at the bottom illustrates the dynamics of the spindle, SPBs, and nuclear mass during mitosis in the *rgf3*ΔN2. Scale bar, 5 $\mu$m.

Given that sister centromere separation (anaphase A) occurs rapidly at the end of Phase II, we next analyzed whether the spindles assembled in *rgf3*ΔN2 cells were competent to segregate chromosomes. Interestingly, the percentage of cells displaying missegregation of the kinetochore marker Mis6 was statistically comparable between both strains (n=70 cells analyzed per strain, Figure S2A). Additionally, cyclin B (Cdc13-GFP), which is targeted for degradation upon proper chromosome bi-orientation, disappeared from both WT and *rgf3*ΔN2 spindles at approximately the same time (Figure S2B). Together, these data indicate that *rgf3*ΔN2 cells effectively separate sister chromatids but fail to properly elongate their mitotic spindles during late anaphase. Finally, we investigated whether premature spindle disassembly could underlie the karyokinesis defect observed in the mutant cells. To test this, we monitored microtubule dynamics in WT and *rgf3*ΔN2 cells expressing Atb2-GFP (α-tubulin). Spindle integrity and kinetics were virtually identical in both strains during Phases I and II (Figure 2D). However, while mitotic spindles in control WT cells remained completely straight throughout nuclear division (Figure 2D, upper panel), anaphase spindles in *rgf3*ΔN2 cells underwent severe buckling (Figure 2D, lower panels). This mechanical buckling frequently caused the spindle to snap, splitting it into two half-spindles and pulling the SPBs back together as the nuclear mass collapsed into a single sphere. This phenomenon visually recapitulates the unusual “fatal attraction” behavior of SPBs characteristic of the *rgf3*ΔN2 mutant. Taken together, these findings demonstrate that although spindles in *rgf3*ΔN2 cells assemble correctly and successfully initiate chromosome segregation, they fail to achieve the critical length required for successful anaphase B due to severe mechanical buckling.

### Dysregulation of Rgf3 inhibits nuclear envelope expansion in anaphase

Fission yeast cells undergo “closed” mitosis, a process wherein mitotic spindle microtubules assemble inside the nucleus and capture chromosomes without dismantling the nuclear envelope (NE). The spindle pole bodies (SPBs) function as the primary mitotic microtubule-organizing centers (MTOCs) and prevent NE deformation during spindle elongation. Consequently, as the elongating spindle encounters the semi-rigid NE, it experiences compressive mechanical stress along its longitudinal axis ^49^. If NE expansion occurs concurrently and parallel to spindle elongation, this compressive stress is minimized, allowing the spindle to remain straight. Conversely, if the nuclear surface area fails to increase, the compressive stress can exceed the spindle’s buckling threshold, triggering severe structural buckling ^4,9,23^.

To determine whether the spindle buckling observed in *rgf3*ΔN2 cells is driven by this mechanical constraint, we monitored NE dynamics and nuclear size throughout mitosis in both wild-type (WT) and mutant strains. Nuclear size was estimated by measuring the maximum cross-sectional area of nuclei in time-lapse micrographs expressing Sur4-mCherry, a marker of the endoplasmic reticulum (ER) and nuclear envelope. In WT control cells, nuclear surface area progressively increased during mitosis, whereas this expansion was significantly suppressed in *rgf3*ΔN2 undergoing SPB missegregation (Figure 3A).

**Figure 3.**
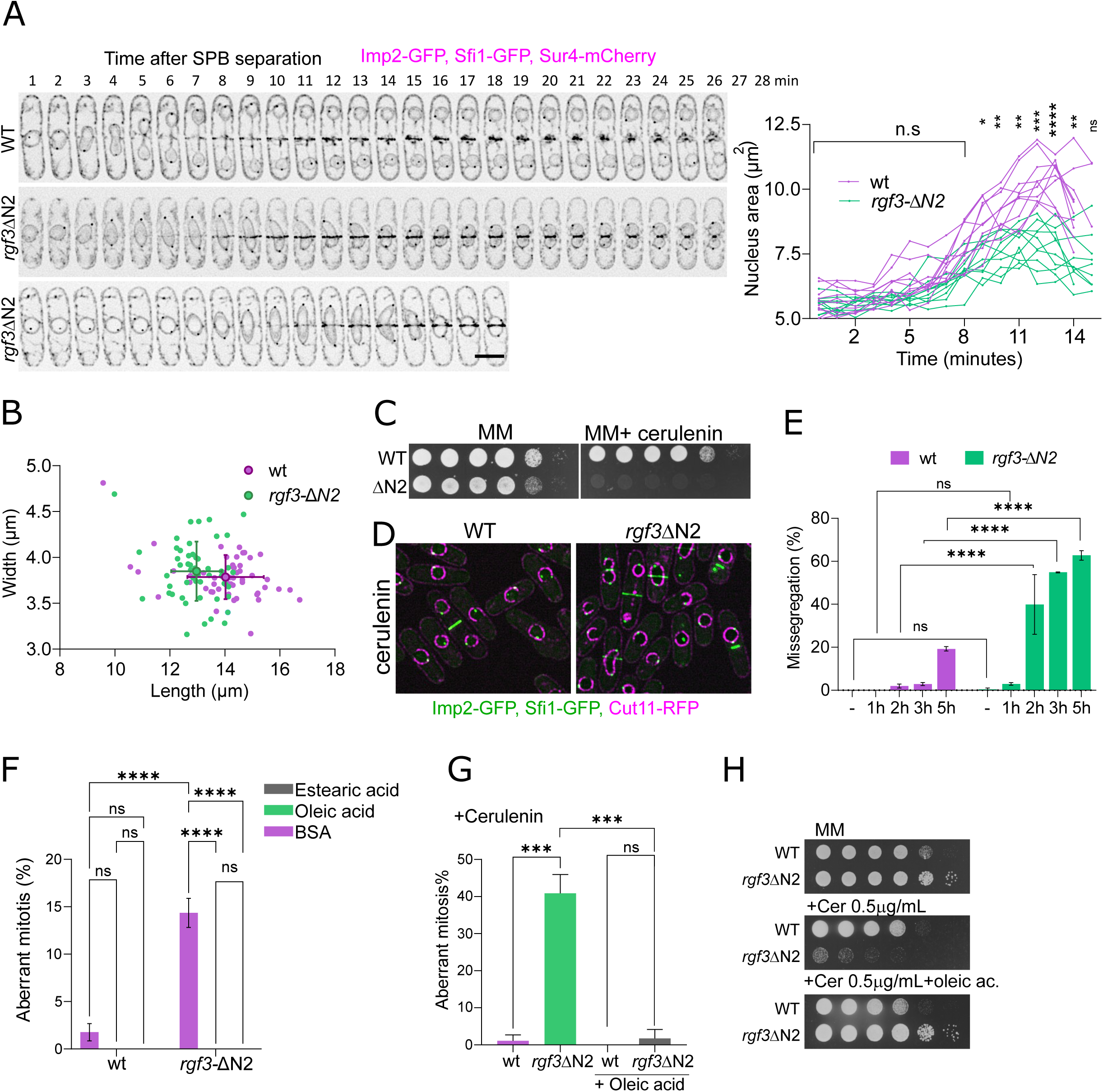
Dysregulation of Rgf3 in anaphase inhibits nuclear envelope expansion. **(A)** Representative time-lapse fluorescence micrographs, shown as inverted grayscale images, of wild-type (WT) and *rgf3*ΔN2 cells expressing Imp2-GFP (CAR marker), Sfi1-GFP (SPB marker), and Sur4-mCherry (ER/nuclear envelope marker). Two representative examples displaying distinct mitotic outcomes are shown for the *rgf3*ΔN2 mutant. Scale bar, 5 µm. (Right panel) Quantitation of the nuclear surface area (µm^2^) over time during nuclear division in WT (n = 10 cells) and *rgf3*ΔN2 mutant (n = 10 cells) strains. Data points represent individual cells; asterisks indicate statistical significance between genotypes at specific time points (* p<0.05; **, p<0.01; ***, p <0.001; ****, p<0.0001: n.s., non-significant; calculated by a two-way ANOVA followed by Sidak’s multiple comparisons test). **(B)** Cell dimensions of septated wild-type (WT) and *rgf3*ΔN2 cells cultured in YES medium at 28°C, determined from DIC micrographs using ImageJ. The average length and width were 12.4 ± 0.67 μm and 4.02 ± 0.37 μm for WT (n = 51), and 10.6 ± 0.72 μm and 3.71 ± 0.25 μm for *rgf3*ΔN2 cells (n=51). **(C)** Functional growth assay in the presence of cerulenin. Strains of WT and *rgf3*ΔN2 were adjusted to an initial O.D._600_ of 2.7. A series of three two-fold serial dilutions followed by two ten-fold serial dilutions were spotted onto minimal media (MM) plates with or without cerulenin (0.5 µg/mL). Colony formation was evaluated after 3–4 days of incubation at 28°C. **(D)** Representative spinning-disk confocal micrographs of WT and *rgf3*ΔN cells expressing Cut11-RFP (nuclear envelope marker), Imp2-GFP (CAR marker), and Sfi1-GFP (SPB marker) following acute cerulenin treatment (5 µM) for 3 h. Scale bar, 7 µm. **(E)** Quantification of the percentage of chromosome missegregation from the cells described in (D). Cells were cultured in MM with or without cerulenin (5 µM) for the indicated time points. **(F)** Percentage of aberrant mitosis in WT and *rgf3*ΔN cells in the absence or presence of 1 mM oleic acid or stearic acid. Cells were grown in MM supplemented with bovine serum albumin (BSA) alone, 1 mM oleic acid plus BSA, or 1 mM stearic acid plus BSA. **(G)** Quantitative analysis of aberrant mitosis percentages. Cells were cultured in MM supplemented with BSA or 1 mM oleic acid for 4 h, followed by the addition of cerulenin (5 µM) and incubation for an additional 3 h prior to imaging. For graphs (E, F, and G), data represent the mean ± SD from three independent experiments (n > 20 cells analyzed per strain per condition). Asterisks denote statistical significance (* p<0.05; **, p<0.01; ***, p <0.001; ****, p<0.0001; n.s., non-significant; calculated by a two-way ANOVA followed by Sidak’s multiple comparisons test). **(H)** Rescue of cerulenin sensitivity by exogenous fatty acids. Strains of WT and *rgf3*ΔN2 were adjusted to an initial O.D._600_ of 2.7. A series of three two-fold serial dilutions followed by two ten-fold serial dilutions were spotted onto control MM plates, MM supplemented with cerulenin (0.5 µg/mL), or MM supplemented with both cerulenin (0.5 µg/mL) and oleic acid (1 mM). Plates were incubated at 28°C until colonies were visible.

Given that nuclear size is proportional to cell size throughout the cell cycle ^50^, we reasoned that the *rgf3*ΔN2 mutant might display an inherent reduction in cell dimensions. Indeed, morphometric measurements at cell division revealed that the average cell length and width were 14.03 ± 1.39 μm and 3.79 ± 0.24 μm in WT cells (n = 51), compared to 12.97 ± 0.92 μm and 3.85 ± 0.32 μm in *rgf3*ΔN2 mutants (n = 51), demonstrating that mutant cells are slightly smaller at division (Figures 3B and S3A, B). Since mitotic nuclear expansion relies strictly on the synthesis of new lipid bilayers, the defect observed in *rgf3*ΔN2 cells could stem from compromised membrane biogenesis, which is tightly linked to nutrient availability and cell size control networks. Lipids serve as the primary structural building blocks for membrane biosynthesis; therefore, we investigated whether limiting *de novo* lipid synthesis would exacerbate the karyokinesis defects of *rgf3*ΔN2 cells. To test this, we utilized cerulenin, an antifungal antibiotic that specifically inhibits *de novo* fatty acid (FA) synthesis ^51^. Restricted membrane availability induced by cerulenin has been shown to cause spindle buckling and aberrant mitosis ^9,52,53^. In a functional spot assay, *rgf3*ΔN2 cells exhibited marked hypersensitivity to cerulenin, displaying severely impaired growth at concentrations (0.5 µg/mL) that had no discernible effect on WT control cells (Figure 3C). Furthermore, acute cerulenin treatment (5 µM) for 1, 2, 3, and 5 h triggered a dramatic, time-dependent increase (>12-fold) in chromosome missegregation frequencies in *rgf3*ΔN2 cells relative to WT controls (Figures 3D and 3E).

To determine whether exogenous lipid supplementation could rescue this phenotype, we supplemented the minimal medium (MM) with either oleic acid (18:1Δ9)—the predominant FA species in *S. pombe* at 20–30°C, accounting for >75% of total cellular FAs—or stearic acid (18:0) ^54,55^. Quantitative fluorescence imaging revealed that supplementation with either FA significantly suppressed the baseline chromosome missegregation defects of *rgf3*ΔN2 cells (Figure 3F). Additionally, exogenous oleic acid supplementation successfully minimized missegregation events in cerulenin-treated *rgf3*ΔN2 cells (Figure 3G) and effectively rescued their growth defects on cerulenin-containing solid media (Figure 3H). Taken together, these findings demonstrate that while the inhibition of FA synthesis severely aggravates the mitotic defects *rgf3*ΔN2 cells ^56^, exogenous FA availability efficiently alleviates them. Given the absolute requirement of fatty acids for membrane biogenesis, our results strongly suggest that the mitotic and mechanical failures observed in *rgf3*ΔN2 cells stem from insufficient lipid availability necessary to sustain nuclear envelope expansion during anaphase elongation.

### The N-terminus of Rgf3 is required for Rho1 activation

To investigate whether Rgf3-mediated Rho1 activation plays a direct role in regulating nuclear envelope (NE) availability, we first re-examined the *in vivo* cellular localization of mNeonGreen-Rgf3 (mNG-Rgf3) and Rho1-sfGFP utilizing Super-Resolution Radial Fluctuations (SRRF) microscopy. Rho1-sfGFP was clearly detected at the plasma membrane (PM), vacuoles, and the endoplasmic reticulum/nuclear envelope (ER/NE) network (Figure 4A). Concurrently, mNG-Rgf3 was found to co-localize at the PM and internal membrane structures, although displaying a substantially weaker basal signal intensity (Figure 4A).

**Figure 4.**
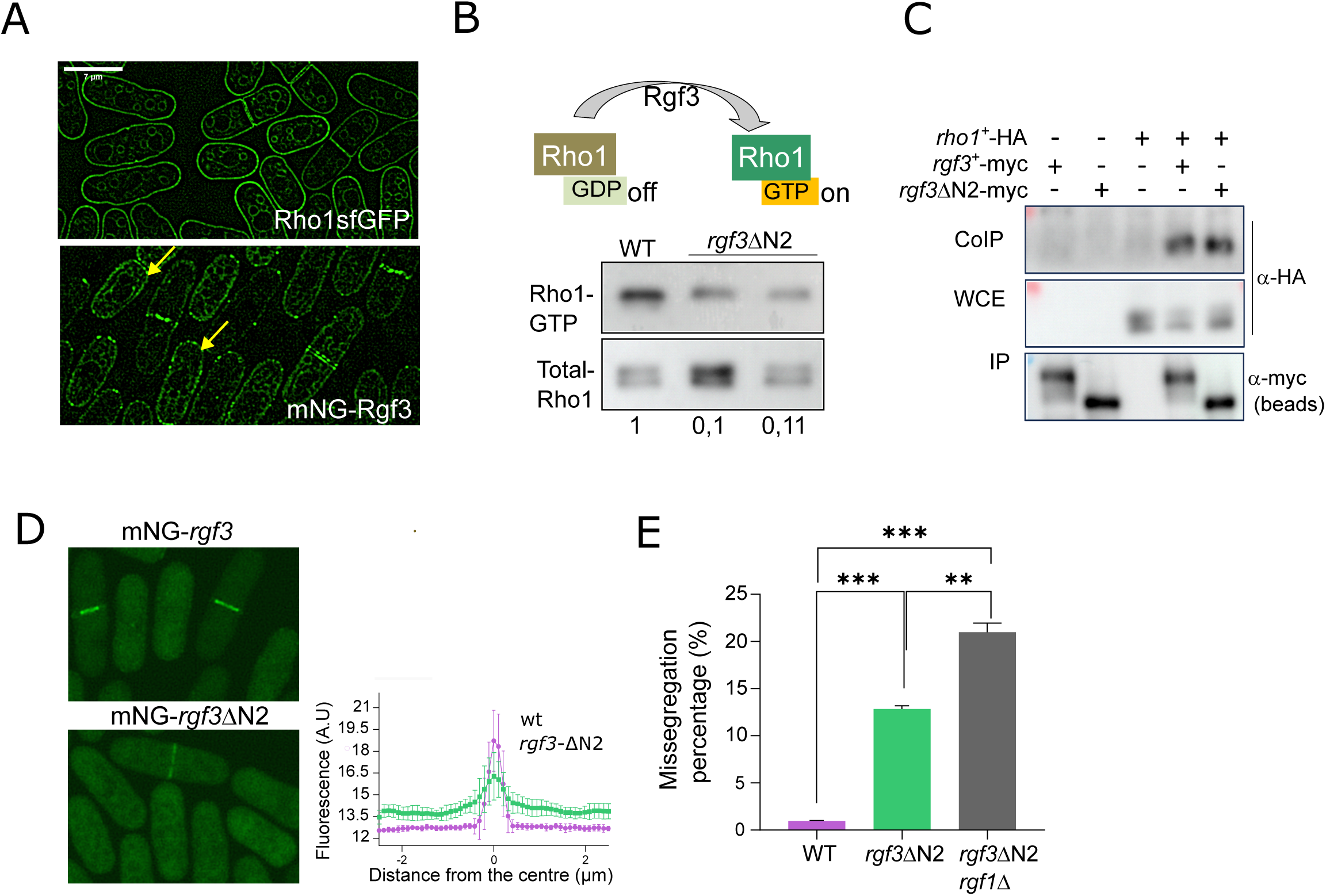
The N-terminus of Rgf3 is required for Rho1 activation. **(A)** Super-Resolution Radial Fluctuation (SRRF) micrographs of wild-type (WT) cells expressing Rho1-sfGFP (integrated at the *leu1* locus) and mNG-Rgf3. Arrows highlight cortical and division-site localization. Scale bar, 7 µm. **(B)** Active Rho1 pull-down assay. Protein extracts from WT (*rgf3*⁺) and *rgf3*ΔN2 cells endogenously expressing HA-tagged Rho1 were subjected to pull-down assays using GST-C21RBD beads, followed by Western blot analysis with an anti-HA antibody to detect active Rho1 (Rho1-GTP). Total Rho1-HA levels were monitored in whole-cell extracts (WCE) as a loading control. Relative units (bottom panel) represent the fold-change in active Rho1 levels compared to the WT control (assigned an arbitrary value of 1.0), representing the mean of two independent experiments. (Top panel) Schematic illustration of the Rho1 GTPase cycle, highlighting activation mediated by GDP/GTP exchange facilitated by the GEF Rgf3. **(C)** Co-immunoprecipitation analysis between Rho1 and mutant Rgf3. Extracts from cells expressing Rgf3-myc, *rgf3*ΔN2-myc, Rho1-HA (controls), or the indicated co-expressing combinations were immunoprecipitated using anti-myc beads and analyzed via Western blot with anti-HA and anti-myc antibodies. Total Rho1-HA levels in WCE were assessed as input control. **(D)** Representative fluorescence images of cells expressing full-length mNG-Rgf3 or mutant mNG-*rgf3*ΔN2. Micrographs are displayed as sum projections of 11-plane - stacks. Scale bar, 5 µm. (Right graph) Fluorescence intensity line scans across the division site for mNG-Rgf3 and mNG-*rgf3*ΔN2 (n= 10 cells per strain). Measurements were performed using a 10-pixel-wide, 5-µm-long line centered on the actomyosin ring. **(E)** Quantification of the SPB missegregation percentage in WT, *rgf3*ΔN2 and *rgf3*ΔN2*rgf1*Δ cells. Data represent the mean ± SD from two independent experiments (n> 20 cells analyzed per strain). Asterisks indicate statistical significance (**, p<0.01; ***, p <0.001; calculated by a one-way ANOVA followed by Tukey’s multiple comparisons test).

We next sought to determine whether the NE expansion defect observed in *rgf3*ΔN2 cells was a direct consequence of impaired Rho1 activation. To test this, we quantified the levels of active, GTP-bound Rho1 in *rgf3*ΔN2 mutant cells via a pull-down assay using a bacterially purified GST-C21RBD (rhotekin-binding domain) probe ^30^. Our results revealed a drastic reduction in active Rho1-GTP levels in cells compared to wild-type (WT) control cells (Figure 4B). This defect in Rho1 activation cannot be attributed to a decrease in Rgf3 protein expression levels, as Western blot analyses demonstrated that the truncated protein is more stable than its full-length WT counterpart (Figure S4A). We reasoned that the truncated protein might either exhibit a reduced binding affinity for Rho1 or display localization defects that prevent proper interaction with its substrate at the division site. To distinguish between these possibilities, we analyzed the physical interaction between mutant Rgf3 and Rho1 from yeast lysates. Intriguingly, endogenously expressed myc-tagged co-immunoprecipitated HA-tagged Rho1 at levels fully comparable to those observed for full-length Rgf3 (Figure 4C), indicating that the N-terminus is dispensable for physical complex formation. However, when we examined the *in vivo* spatial distribution of mNG-*rgf3*ΔN2, the mutant protein exhibited significantly weaker accumulation at the actomyosin ring and division site, remaining predominantly cytoplasmic compared to full-length mNG-Rgf3 (Figure 4D). Thus, while both full-length Rgf3 and the mutant retain the capacity to bind Rho1, only the truncation results in severely reduced Rho1-GTP levels, underscoring the critical requirement of the N-terminal domain for promoting proper GDP/GTP exchange.

Because Rho1 is an essential GTPase and available conditional mutants exhibit severe pleiotropic defects ^57^, directly assessing the impact of a total loss of Rho1 function on the spindle buckling phenotype remains technically challenging. To circumvent this and validate our model, we evaluated a genetic interaction by deleting *rgf1*, which encodes the other principal Rho1 GEF in fission yeast vegetative cells. Remarkably, in the double mutant strain (which severely compromises overall Rho1 activation), the chromosome missegregation rate significantly escalated to approximately 22%, compared to the 12% observed in the single *rgf3*ΔN2 mutant (Figure 4E). Taken together, these genetic and biochemical lines of evidence indicate that Rgf3 functions upstream of Rho1 to drive the lipid-mediated nuclear envelope expansion required for successful chromosome segregation during anaphase.

### The *rgf3-*Δ*N2* mutant limits the availability of the NM in lipin pathway mutants

The growing evidence that lipid biogenesis plays a crucial role in nuclear envelope (NE) enlargement during mitosis ^9,16,56,58^ prompted us to investigate whether Rho1 and its GEF Rgf3 interact directly with lipids. Utilizing a protein–lipid overlay assay, we found that recombinant Rgf3, expressed as a GST-fusion protein in *E. coli*, binds directly to several negatively charged lipids, including phosphatidic acid (PA), phosphatidylinositol phosphates (PtdIns-4-P and PtdIns-4,5-P2), cardiolipin (CL), and sulfatides (Figure 5A). In the same assay, Rho1 specifically interacted with PA and phosphatidylserine (PS) (Figure 5A). Given that Rgf3 localizes to the NE/ER boundaries and that PA and phosphatidylinositol phosphates are major ER components ^59,60^, Rgf3-mediated activation of Rho1 might regulate local PA availability or downstream lipid derivatives. PA is a central metabolic precursor for both glycerophospholipid (GPL) biosynthesis and triacylglycerol (TAG) storage, and its precise regulation is critical for successful NE expansion during mitosis ^23,61,62^ (Figure 5B). PA is synthesized via the stepwise acylation of glycerol-3-phosphate and can subsequently be dephosphorylated to diacylglycerol (DAG) or converted into CDP-DAG to drive GPL synthesis ^15^. This DAG-to-PA interconversion is catalyzed antagonistically by the diacylglycerol kinase Dgk1 and the lipin phosphatase Ned1, respectively. Ned1 activity is negatively regulated by CDK1 (Cdc2) phosphorylation and positively sustained by the Nem1/Spo7 phosphatase complex (Figure 5B) ^16^. Consistently, loss of lipin, Nem1, or Spo7 causes hyperproliferation of nuclear and peripheral ER membranes ^16^ and reduces lipid droplet formation. These observations led us to hypothesize that the membrane restriction defects associated with the *rgf3*ΔN2 mutant could counteract the membrane overgrowth phenotypes characteristic of lipin pathway mutants. Indeed, introducing the *rgf3*ΔN2 mutation partially restored a normal, quasi-spherical nuclear shape in *nem1*Δ cells, successfully reversing their characteristic flaccid NE phenotype (Figure 5C). In agreement with this, morphometric quantification revealed that single *rgf3*ΔN2 cells contained a slightly higher number of lipid droplets (8.7 ± 2.2; mean ± SD) relative to wild-type (WT) control cells (6.62 ± 1.56). Moreover, the *rgf3*ΔN2 *nem1*Δ double mutant partially suppressed the depleted lipid-droplet phenotype of *nem1*Δ single mutants (6.17 ± 2.23 versus 4.05 ± 1.63, respectively; Figure S5A) and significantly mitigated their slow-growth phenotype on minimal medium (MM) plates (Figure 5D).

**Figure 5.**
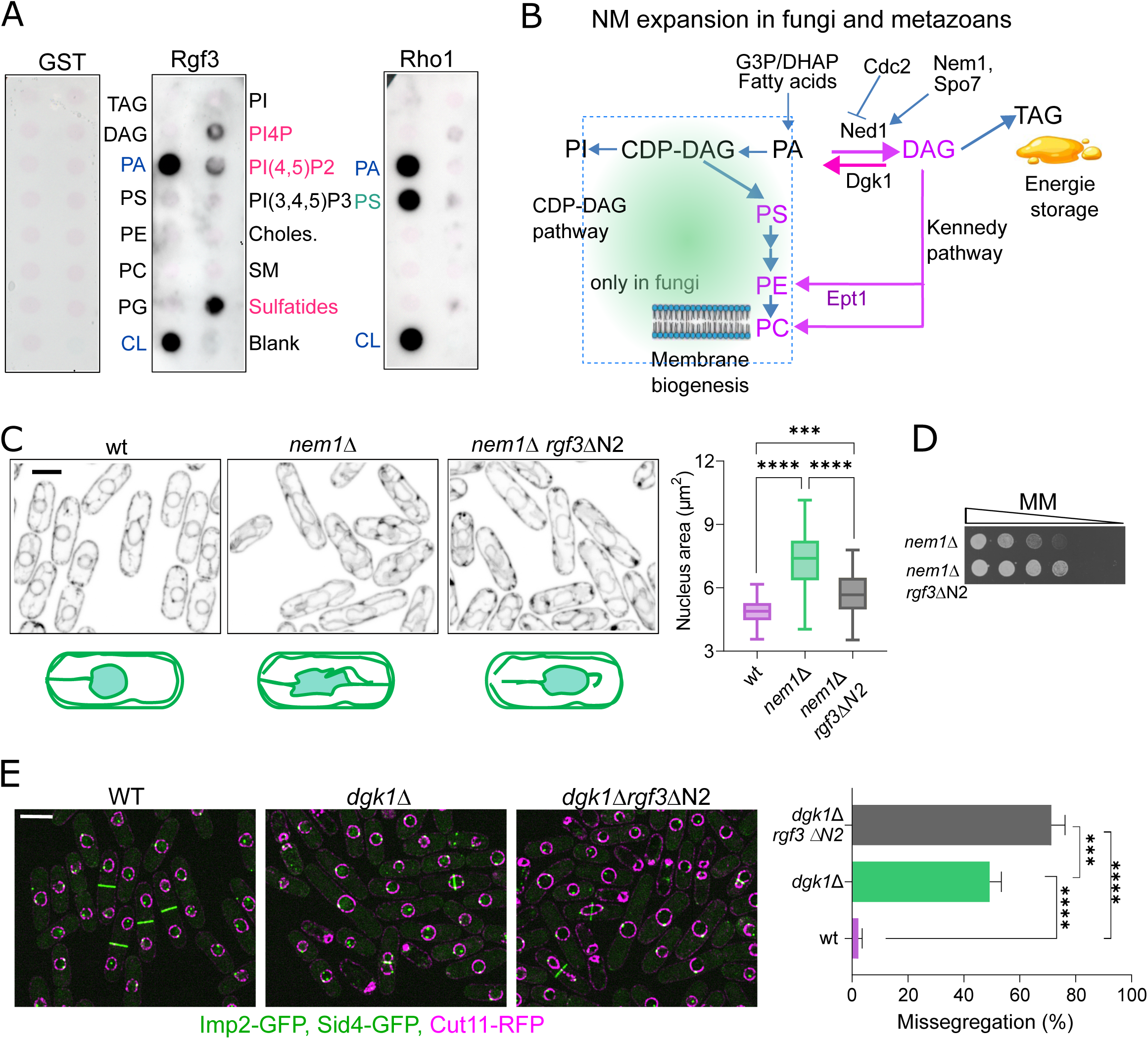
The rgf3ΔN2 mutant restricts NM availability in lipin pathway mutants. **(A)** Protein-lipid overlay assay evaluating GST-Rgf3 and GST-Rho1 binding specificities. Membrane lipid strips were incubated with 1 µg/ml of purified GST (control), Rgf3-GST, or Rho1-GST, and macromolecular interactions were detected using an anti-GST antibody. Lipids significantly bound by Rgf3-GST are highlighted in pink, those bound by Rho1-GST in green, and shared target lipids in blue. Both immunoblots were processed under identical experimental conditions. **(B)** Schematic model illustrating phosphatidate (PA) and diacylglycerol (DAG) metabolic flux during nuclear envelope (NE) expansion in *S. pombe* (adapted from reference ^23^ and results from this study). PA serves as a critical precursor for major glycerophospholipids (GPLs) required for membrane biogenesis, whereas triacylglycerol (TAG) is funneled into lipid droplets for energy storage. During mitosis, inactivation of the Ned1 lipin by CDK1 (Cdc2), coupled with diacylglycerol kinase Dgk1 activity, channels DAG toward GPL biosynthesis. In the absence of Dgk1, compromised DAG-to-PA conversion restricts nuclear membrane availability. Alternatively, cells rely on the Kennedy pathway (KP) for GPL production. In *ept1* mutants, which are unable to transfer activated choline or ethanolamine to DAG, the KP is inactive, resulting in restricted NE expansion and a characteristic “cut” phenotype. Abbreviations: CDP-DAG, CDP-diacylglycerol; G3P, glycerol-3-phosphate; DHAP, dihydroxyacetone phosphate; PC, phosphatidylcholine; PE, phosphatidylethanolamine; PI, phosphatidylinositol; PS, phosphatidylserine. The synthesis of PE and PC from lyso phosphatidylethanolamine (LyPE) and lyso phosphatidylcholine (LyPC), respectively, is not shown in the figure. **(C)** Representative fluorescence micrographs of cells expressing Sur4-mCherry in WT, *nem1*Δ, and *nem1*Δ*rgf3*ΔN2 backgrounds. Images represent maximum-intensity projections of six Z-sections (0.5 µm step-size). (Right panel) Quantitation of the nuclear surface area (µm^2^) during nuclear division in WT (n = 50), *nem1*Δ (n = 50), and *nem1*Δ*rgf3*ΔN2 (n = 50) cells. Box plots define the interquartile range (IQR); whiskers represent Tukey’s range, and the horizontal line inside the box indicates the median value. Statistical significance was calculated using a one-way ANOVA followed by Šidák’s multiple comparisons test (***, p <0.001; ****, p<0.0001). Scale bar, 5 µm. **(D)** Functional growth assay of *nem1*Δ and *nem1*Δ *rgf3*ΔN2 strains. Cells were adjusted to an initial of 2.7, and a series of three two-fold serial dilutions followed by two ten-fold serial dilutions were spotted onto minimal media (MM) solid plates. Colony formation was assessed after 3–4 days of incubation at 28°C **(E)** (Left panels) Representative fluorescence micrographs of WT, *dgk1*Δ, and *rgf3*ΔN2 *dgk1*Δ mutant cells expressing Cut11-RFP (nuclear envelope marker), Imp2-GFP (CAR marker), and Sid4-GFP (SPB marker). Scale bar, 7 µm. (Right panel) Quantitative analysis of the percentage of cells displaying aberrant mitosis/aberrant nuclear size in the indicated strains. Data represent the mean ± SD from three independent experiments (n>20 cells analyzed per strain). Asterisks indicate statistical significance (***, p <0.001; ****, p<0.0001; calculated by a one-way ANOVA followed by Tukey’s multiple comparisons test).

One potential mechanism is that Rgf3/Rho1 negatively regulates the Nem1–Spo7 complex, meaning that the loss of the Rgf3 N-terminus would indirectly enhance Ned1 activation via dephosphorylation. However, an analysis of the phosphorylation status of Ned1 in mitotic cells released from a hydroxyurea (HU) block revealed no significant differences in Ned1 electrophoretic mobility between WT and *rgf3*ΔN2 cells (Figure S5C), ruling out a direct regulatory effect on Ned1 phosphorylation. We next asked whether deletion of Dgk1 kinase, which catalyzes the conversion of DAG to PA and thus directly antagonizes lipin activity (Figure 5B), would conversely exacerbate the defects of the *rgf3*ΔN2 mutant. Loss of Dgk1 was recently shown to rescue the NE and ER expansion phenotypes of lipin pathway mutants in *S. pombe* ^23^. In our experiments, over 45% of single *dgk1*Δ cells displayed abnormal, bow-shaped nuclear intermediates during anaphase, accompanied by a severe failure in NE expansion. Strikingly, the *rgf3*ΔN2 *dgk1*Δ double mutant exhibited a strong synthetic-negative phenotype: ∼75% of cells displayed severe chromosome “cut” or aberrant mitosis phenotypes, compared with 45% in *dgk1*Δ single mutants and negligible levels in WT cells (Figure 5E). Consistently, growth of the double mutant was markedly more impaired than that of the *dgk1*Δ single mutant in a drop dilution assay (Figure S5B).

Together with our previous findings, these results demonstrate that while the *rgf3*ΔN2 defect can alleviate the excess NE phenotype caused by lipin pathway inactivation, it synergistically worsens the defects of PA-depleted cells (*dgk1*Δ). This strongly reinforces the model that Rgf3/Rho1 activity acts to promote NE expansion. Importantly, the strong synthetic interaction between *rgf3*ΔN2 and *dgk1*Δ suggests that Rgf3/Rho1 functions in a pathway parallel to Dgk1 (synthesis of PA from DAG) to support proper nuclear envelope remodeling and expansion during mitotic karyokinesis.

### Rgf3 and Rho1 regulate glycerophospholipid production necessary for nuclear envelope expansion

To examine the role of Rgf3/Rho1 in nuclear envelope (NE) expansion, we quantified the major lipid species—glycerophospholipids (GPLs), DAG, and triacylglycerols (TAG)—in the *rgf3*ΔN2 mutant relative to wild-type (WT) cells. Quantitative lipidomic analysis revealed that the mutant displayed a small but statistically significant reduction in phosphatidylcholine (PC) and lysophosphatidylcholine (lysoPC) (Figure 6A), whereas overall cellular DAG and TAG levels remained statistically unchanged (Figure S6A). Given that the *rgf3*ΔN2 mutation synergistically exacerbates the NE expansion defects observed in a *dgk1*Δ background, we hypothesized that reduced Rho1 activation in *rgf3*ΔN2 cells compromises PC synthesis through a route parallel or alternative to the *de novo* CDP–DAG pathway. As observed in higher eukaryotes, DAG serves not only as a structural precursor for TAG synthesis but also as a critical substrate for the production of phosphatidylethanolamine (PE) and PC via the CDP–ethanolamine and CDP–choline branches of the Kennedy pathway (KP) ^14,63^ (Figure 6B).

**Figure 6.**
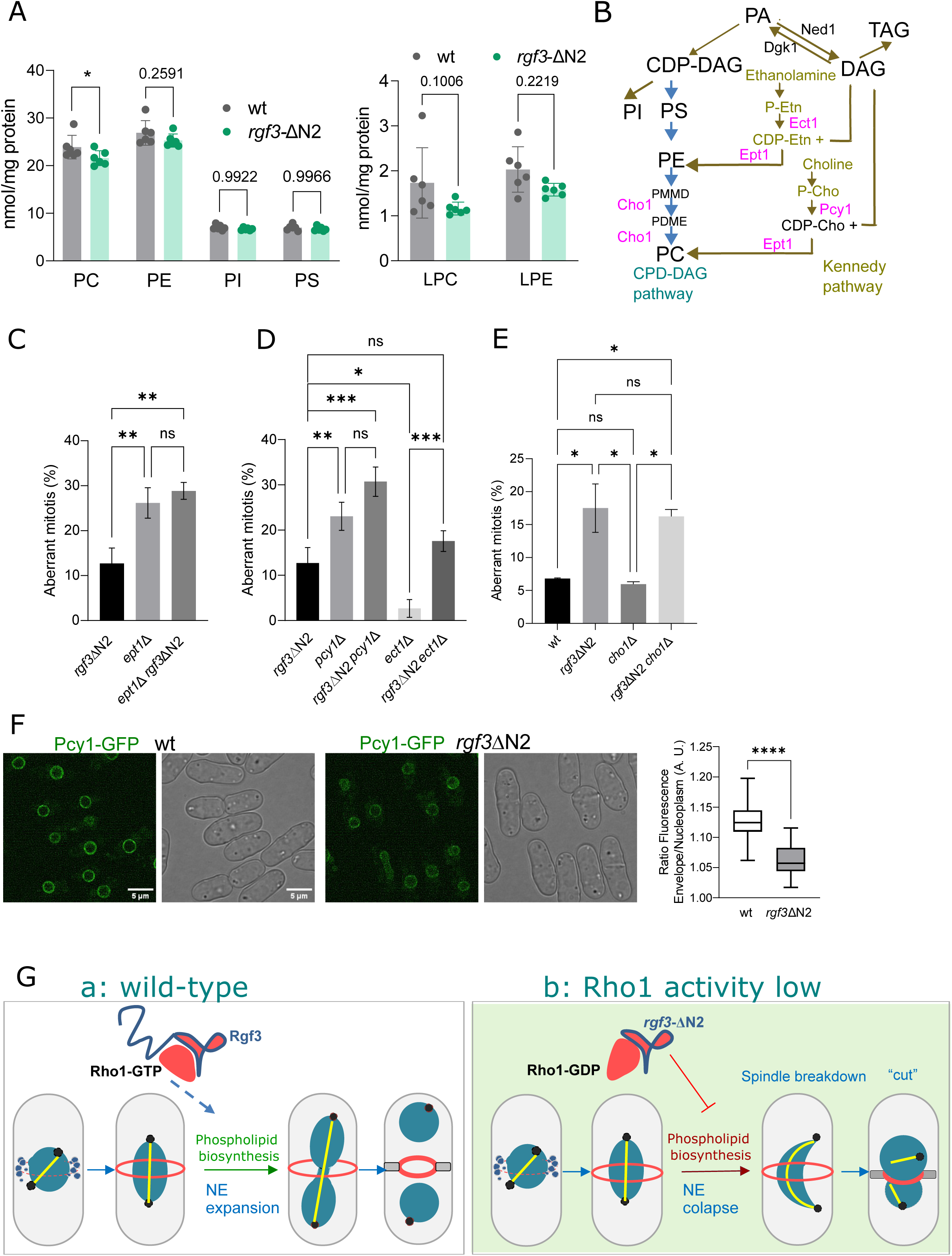
Lipidomic analysis and genetic interactions with the phospholipid biosynthesis pathways. **(A)** Quantitative lipidomic analysis of wild-type (WT) and *rgf3*ΔN2 cells. (Left graph) Relative abundance (nmol/mg protein) of the four major glycerophospholipid (GPL) classes: phosphatidylcholine (PC), phosphatidylethanolamine (PE), phosphatidylinositol (PI), and phosphatidylserine (PS). (Right graph) Relative abundance of lysophosphatidylcholine (LPC) and lysophosphatidylethanolamine (LPE). Data represent the mean ±SD from n = 6 biological replicates. Asterisks indicate statistical significance compared to WT cells calculated by a two-tailed Student’s *t*-test assuming equal variance (*, p < 0.05). **(B)** Schematic diagram of the major phospholipid synthesis pathways in *S. pombe*. Phosphatidic acid (PA) serves as the central metabolic precursor for phospholipid (PL) biosynthesis, channeling into cytidine diphosphate-diacylglycerol (CDP-DAG) production. ER-localized CDP-DAG drives the synthesis of PI and PS. PS undergoes decarboxylation to form PE, which is sequentially methylated three times to yield PC, the most abundant PL in *S. pombe*. Alternatively, PE and PC can be synthesized from exogenous ethanolamine and choline via the Kennedy pathway. Enzymes catalyzing individual metabolic steps are indicated in pink. Abbreviations: CDP-DAG, CDP-diacylglycerol; DAG, diacylglycerol; TAG, triacylglycerol; Cho, choline; Etn, ethanolamine; P-Cho, phosphocholine; P-Etn, phosphoethanolamine; CDP-Cho, CDP-choline; CDP-Etn, CDP-ethanolamine; PMMD, phosphatidyldimethylethanolamine; PDME, phosphatidylmonomethylethanolamine. **(C)** Quantification of the percentage of aberrant mitosis in *rgf3*Δ*N2*, *ept1*Δ and *rgf3*Δ*N2 ept1*Δ mutant cells. Data represent the mean ± SD from three independent experiments (n > 20 cells analyzed per strain). Statistical significance is indicated by asterisks (**, p < 0.01; n.s., non-significant; determined by a one-way ANOVA followed by Tukey’s multiple comparisons test). **(D)** Quantification of the percentage of aberrant mitosis in *rgf3*Δ*N2*, *pcy1*Δ, *ect1*Δ, *rgf3*Δ*N2 pcy1*Δ and *rgf3*Δ*N2 ect1*Δ cells. Data represent the mean ± SD from three independent experiments (n > 20 cells analyzed per strain). Asterisks denote statistical significance (*, p< 0.05; **, p< 0.01; ***, p< 0.001; n.s., non-significant; calculated via one-way ANOVA followed by Tukey’s test). **(E)** Quantification of the percentage of aberrant mitosis in WT, *rgf3*Δ*N2*, *cho1*Δ and *rgf3*Δ*N2 cho1*Δ cells. Data represent the mean ± SD from three independent experiments (n > 20 cells analyzed per strain). Significance was assessed using a one-way ANOVA followed by Tukey’s test (*, p < 0.05; n.s., non-significant). **(F)** Subcellular distribution and quantitation of Pcy1-GFP. Fluorescence intensities at the nuclear envelope (NE) and nucleoplasm were measured, and the resulting NE-to-nucleoplasm fluorescence ratio was plotted for WT (n = 50) and *rgf3*ΔN2 (n = 50) cells. Box plots display the interquartile range (IQR); whiskers represent Tukey’s range, and the horizontal line within each box indicates the median value. Statistical significance was determined using a Student’s t-test (****, p < 0.0001). **(G)** Proposed mechanistic model for Rho1-dependent nuclear envelope expansion. (Panel a) In WT cells, Rgf3-mediated activation of Rho1 drives a coordinated burst of phospholipid biosynthesis, facilitating proper nuclear envelope expansion during anaphase. (Panel b) In mutant cells lacking the N-terminal region of Rgf3 *rgf3*ΔN2, compromised Rho1 activity perturbs the Kennedy pathway for PC biosynthesis, leading to restricted membrane availability, NE collapse, and spindle breakdown, resulting in a characteristic “cut” phenotype.

To assess the specific contribution of PC and PE derived from exogenous choline (Cho) and ethanolamine (Etn) precursors to mitotic NE expansion, we analyzed the frequency of mitotic defects in various KP null mutants (Figure 6B). In rich medium, approximately 26% of dividing cells lacking the diacylglycerol cholinephosphotransferase/ethanolaminephosphotransferase (Ept1, which catalyzes the final condensation step of the KP) and about 22% of cells lacking the choline-phosphate cytidylyltransferase (Pcy1, defective in the CDP–choline branch) exhibited severe aberrant mitosis phenotypes (Figure 6C and 6D). By contrast, cells lacking the ethanolamine-phosphate cytidylyltransferase (Ect1, defective in the CDP–ethanolamine branch) behaved indistinguishably from WT controls (Figure 6D). These findings demonstrate that KP activity is strictly required for proper NE expansion and suggest that the PC-producing CDP–choline branch plays a more critical role during mitosis than the PE-producing CDP–ethanolamine branch. This is consistent with the physiological fact that PC is the most abundant phospholipid in eukaryotic membranes and that its sustained synthesis is essential for cellular processes involving rapid membrane surface growth ^64,65^.

We next took advantage of genetic interactions to determine whether the mitotic defects observed in KP mutants were epistatic to or influenced by reduced Rho1 activity. Strikingly, the percentage of aberrant mitosis in the *rgf3*ΔN2 *ept1*Δ and *rgf3*ΔN2 *pcy1*Δ double mutants was fully comparable to that observed in their respective *ept1*Δ and *pcy1*Δ single mutants (Figure 6C and 6D), indicating that elimination of the Rgf3 N-terminus yields no cumulative defects in the absence of a functional KP. This suggests a formal epistatic relationship between the Rgf3/Rho1 module and KP components. Consistently, the *rgf3*ΔN2 *ect1*Δ double mutant (defective in the non-essential CDP–ethanolamine branch) was phenotypically indistinguishable from the *rgf3*ΔN2 single mutant (Figure 6D).

We next examined whether blocking de novo GPL synthesis through the CDP–DAG pathway (also known as the methylation pathway) would conversely affect NE expansion. This route represents the primary source of cellular PC when exogenous Cho is restricted in the growth medium. Interestingly, mutants lacking the CDP–DAG pathway enzyme Cho1 did not exhibit significant mitotic segregation defects (Figure 6E). Likewise, the aberrant mitosis rates in the *rgf3*ΔN2 *cho1*Δ double mutant closely resembled those of the *rgf3*ΔN2 single mutant (Figure 6E), supporting the notion that in rich medium containing Cho and Etn, the KP alone is sufficient to sustain the lipid biosynthesis required for proper NME. In line with these observations, mutants compromised in the CDP–DAG pathway (e.g., *cho1*Δ, *cho2*Δ) behave as strict choline auxotrophs on minimal media, whereas *rgf3*ΔN2 cells grow robustly in minimal medium (MM) lacking both Cho and Etn (Figure S6B), further validating a fully functional *de novo*

CDP–DAG pathway in the mutant background. Furthermore, while exogenous Cho supplementation efficiently suppresses the “cut” phenotype of *dgk1*Δ cells ^23^, it failed to elicit a statistically significant rescue of the mitotic NE expansion defects in *rgf3*ΔN2 cells (Figure S6C). Taken together, these lines of genetic evidence suggest that Rgf3/Rho1 activity contributes to mitotic NE expansion primarily through the KP—most likely by promoting efficient PC synthesis in the presence of exogenous precursors—with the CDP–DAG methylation pathway playing a minor or secondary role under these growth conditions.

### Rgf3/Rho1 signaling regulates GFP-Pcy1 association with the nuclear membrane

Next, we sought to elucidate the functional basis of the mechanistic relationship between PC synthesis through the Kennedy pathway (KP) and impaired Rho1 activation. In most eukaryotes, the CDP–choline branch of the KP represents the major route for cellular PC biosynthesis (Figure 6B) ^66^. This metabolic pathway initiates with the phosphorylation of exogenous choline by choline kinase (Eki1, systematic ID: SPAC13G7.12c), generating phosphocholine. In the subsequent rate-limiting enzymatic step, phosphocholine and CTP are converted to CDP–choline by the CTP:phosphocholine cytidylyltransferase, encoded by the *pcy1*⁺ gene (systematic ID: SPCC1827.02c). Finally, the cholinephosphotransferase Ept1 catalyzes the transfer of phosphocholine from CDP–choline to diacylglycerol (DAG), resulting in the structural formation of PC ^23^ (Figure 6B).

To investigate whether reduced Rho1 activity directly influences the spatial behavior or stability of these key KP enzymes, we analyzed the subcellular localization of GFP-Pcy1 and Ept1-GFP in both WT and *rgf3*ΔN2 mutant cells. *S. pombe* Pcy1 is homologous to *S. cerevisiae* Pct1 and mammalian PCYT1A/PCYT1B (CCTα and CCTβ). These cytidylyltransferases are classic amphitropic enzymes whose physical association with cellular membranes is closely coupled to enhanced catalytic activity ^67^. In budding yeast, Pct1 localizes to both the inner nuclear membrane (INM) and the nucleoplasm, and its active recruitment to the INM is heavily stimulated under conditions of elevated PC demand ^68^. Similarly, in *S. pombe*, N-terminally tagged GFP-Pcy1 re-localizes from the nuclear envelope (NE) to the nucleoplasm during stationary phase, a physiological state where membrane biogenesis declines and storage lipid synthesis predominates (Figure S6D), reinforcing the notion that membrane association is required for full enzymatic activation. We found that GFP–Pcy1 localized to both the NE and the nucleoplasm in WT and *rgf3*ΔN2 cells. However, in *rgf3*ΔN2 mutant cells, GFP–Pcy1 accumulation at the NE was visibly reduced, while its signal within the nucleoplasm became more prominent. Quantitative morphometric analysis confirmed a statistically significant decrease in the NE-to-nucleoplasm fluorescence ratio of GFP-Pcy1 in *rgf3*ΔN2 compared with WT cells (Figure 6F), whereas total Pcy1 protein expression levels remained fully comparable between both strains, as confirmed by immunoblotting (Figure S6E). These results demonstrate that the proper enrichment of Pcy1 at its site of action is compromised under conditions of reduced Rho1 signaling. Conversely, no spatial differences were observed in the subcellular localization of Ept1–GFP between the strains (Figure S6F). Collectively, these findings indicate that Rho1 activity is strictly required for the proper membrane association and activation of Pcy1, and consequently, for efficient PC synthesis during mitotic NE expansion (NME). Active Rho1-GTP may generate or sustain the specific lipid environment necessary for recruiting or stabilizing Pcy1 at active sites of membrane growth. Given that both the Kennedy pathway and Rho1 (RhoA in mammals) are evolutionarily conserved across eukaryotes, our findings uncover a previously unrecognized mechanism through which Rho GTPases modulate phospholipid biosynthesis. More broadly, this study highlights the critical role of Rho proteins—traditionally viewed as master regulators of cell polarity and cortical morphogenesis—in coordinating active membrane biogenesis with cell cycle progression, particularly during mitotic exit.

## Discussion

### Rgf3 is involved in NM expansion at anaphase

We present compelling evidence that the activation of Rho1 by Rgf3 is required to achieve the NE expansion necessary to complete mitosis exit safely. During anaphase, before the actomyosin ring constricts and septum division progresses, cells must expand the NE in strict coordination with intranuclear mitotic spindle elongation and chromosome segregation ^62^. These events are directed by surveillance mechanisms (checkpoints) that tightly monitor both their order and fidelity ^42,69^. However, it remains unknown how the precise timing of nuclear envelope expansion (NME) is determined. Is it merely a passive consequence of physical spindle elongation, or is it an independently regulated process controlled by the cell cycle machinery? Moreover, how is contractile actomyosin ring (CAR) constriction coordinated with NME to prevent the constricting ring and septum from physically severing the dividing nuclei? Our results suggest that, in organisms undergoing closed mitosis, nuclear envelope expansion occurs independently of spindle elongation ^70^. Instead, it takes place in response to specific cell cycle cues that target phospholipids metabolism to supply the main lipid components required for NE growth^9,53,58^.

Approximately 15% of dividing *rgf3*ΔN2 mutant cells, which lack the regulatory N-terminus of Rgf3, display prominent defects in SPB separation. This defect frequently leads to the formation of asymmetrical anucleated and uninucleated cells with nuclei of different sizes (Figure 1D). Notably, *rgf3*ΔN2 cells assemble functional mitotic spindles capable of segregating chromosomes (Figures S2), but these spindles are unable to achieve the final length necessary to fully separate the two daughter nuclei because they undergo severe bending and buckling during anaphase B (Figure 2). This phenotypic outcome can be explained within the context of “closed” mitosis, wherein the nucleus must divide without losing its nucleocytoplasmic compartmentalization ^3,5,62^. In *S. pombe* mitosis, the mother nucleus sequentially transforms into a “rugby ball” and further elongates into a characteristic “dumbbell structure” that ultimately gives rise to a pair of sphere-shaped daughter nuclei. The whole process occurs rapidly in less than 5 minutes (Figure 3A). To accommodate such dramatic spindle elongation, the nuclear surface area must increase by approximately 33% while the overall nuclear volume remains constant ^2,9,56,71^.

In the *rgf3*ΔN2 mutant, NE extension fails to occur in parallel with spindle elongation; consequently, the resulting compressive stress ultimately exceeds the buckling threshold of the microtubule spindle, causing it to bend and collapse. Several independent lines of evidence support this mechanical model: (i) the typical mitotic increase in nuclear size exhibited by wild-type cells is largely abolished in *rgf3*ΔN2 cells displaying the “cut” phenotype (Figure 3A); (ii) *rgf3-*ΔN2 cells are unable to grow in the presence of ceruleninan inhibitor of *de novo* fatty acid synthesis that limits membrane lipid availability- and acute cerulenin treatment triggers a dramatic 10-fold increase in the percentage of aberrant mitosis in asynchronous cultures; and (iii) The *rgf3-*ΔN2 mutation synthetically restricts NE availability when combined with lipin pathway mutants (Figure 5C) ^16,23^. Taken together, these results demonstrate that Rgf3 plays a pivotal role in regulating the membrane availability required for mitotic NME.

This physiological function critically depends on the N-terminal region of Rgf3, which governs its cell cycle-dependent subcellular localization and mediates Rho1 activation (Figure 4D and 4B). Interestingly, this N-terminus is a structurally flexible, low-complexity domain exhibiting classic features of intrinsically disordered regions (IDRs). Similar to other characterized IDRs, the N-terminus of Rgf3 is highly enriched in polar and proline residues while being markedly depleted of hydrophobic amino acids—biophysical properties that prevent the domain from adopting a stably folded tertiary conformation. Due to their structural plasticity, IDRs can transiently adopt multiple conformations depending on their specific binding partners, thereby facilitating highly dynamic interactions with a wide array of downstream targets ^72,73^. Remarkably, within the N-terminus of Rgf3, approximately 29% of the amino acid residues are predicted targets for serine/threonine phosphorylation, forming potential consensus motifs for numerous cell cycle kinases ^74^. This strongly suggests that precise phosphorylation or dephosphorylation events within this disordered domain may act as a critical regulatory switch to fine-tune Rho1 activation, thereby driving efficient nuclear envelope expansion during mitotic exit.

### Rgf3-mediated Rho1 activation is required to maintain phospholipid homeostasis during mitosis

A key question arising from our findings is how mutations in Rgf3 that impair Rho1 activation lead to severe defects in nuclear envelope expansion (NME). From a biophysical perspective, the arched-spindle phenotype implies the presence of counteracting, anti-elongation forces acting on the NE. In wild-type cells, deformation of the nucleus from its default spherical shape during anaphase is tightly accommodated by structural phospholipid synthesis—through head-group addition and membrane remodeling—which supplies fresh lipid material while preserving NE integrity and fluidity. Our results indicate that the *rgf3* mutants fail to properly expand the NE because they are unable to upregulate phospholipid synthesis during mitosis. These observations can be integrated into a model describing the regulatory network that controls glycerophospholipid (GLP) synthesis (Figure 5B). In fungi, the major route for GLP production proceeds through the conversion of phosphatidic acid (PA) to CDP–DAG, via the *de novo* CDP–DAG pathway ^14,15^. Consistent with this framework, mutants defective in the lipin pathway exhibit excessive nuclear envelope outgrowths, or “nuclear flares” ^16^, whereas *dgk1*Δ cells—which are unable to regenerate PA from DAG—display the opposite phenotype, failing to expand the NE ^23^. The mitotic phenotype of *rgf3*ΔN2 cells closely resembles that of *dgk1*Δ, initially suggesting that GLP synthesis through the CDP–DAG pathway might be compromised. However, several lines of evidence argue against this interpretation. First, lipin (Ned1) activity appears unaltered in *rgf3*ΔN2 cells, as indicated by its normal mitotic phosphorylation pattern (Figure S5C). Second, null mutants affecting late steps of the CDP–DAG pathway do not display chromosome “cut” phenotypes. Finally, the synergistically exacerbated mitotic defects observed in the *dgk1*Δ *rgf3*ΔN2 double mutant demonstrate that Rgf3 and Dgk1 act in distinct, parallel pathways that collectively promote NE expansion and lipid homeostasis (Figure 5E). Our analyses further suggest that the reduction in PC and LPC levels observed in *rgf3*ΔN2 cells (Figure 6A) arises from specific defects in the CDP–choline branch of the Kennedy pathway (KP). This pathway constitutes the principal route for PC and phosphatidylethanolamine (PE) biosynthesis in metazoans, where DAG serves as the essential lipid substrate for PC and PE formation through the enzymatic transfer of CDP–choline and CDP–ethanolamine, respectively (Figure 6B). In fungi, the KP is primarily active under conditions where choline and ethanolamine are available in the environment, and the presence of exogenous choline feedback-represses the alternative CDP–DAG methylation pathway ^75^. PC represents more than 50% of total cellular phospholipids, and its sustained synthesis is essential for cellular processes involving rapid membrane surface growth, including cell division and organelle expansion. We found that *S. pombe* mutants anchoring the PC branch of the KP (*pcy1*Δ and *ept1*Δ) display severe nuclear division defects (Figure 6C and 6D), reinforcing the critical role of the KP in NME previously proposed by Oliferenkós laboratory ^23^. Importantly, localized Rho1 activity ensures the proper recruitment of the rate-limiting enzyme Pcy1 to the nuclear envelope, facilitating the efficient conversion of phosphocholine and DAG into PC. This local lipid synthesis, in turn, supports membrane biogenesis and NE expansion as the microtubule spindle elongates. In the absence of Rgf3-mediated activation, reduced Rho1 activity leads to GFP-Pcy1 mislocalization, decreased PC production, and insufficient membrane supply, ultimately resulting in NE deformation and spindle buckling (Figure 6F).

CTP:phosphocholine cytidylyltransferases (CCTs) contain a conserved C-terminal inhibitory “M” domain that represses enzymatic activity through an intramolecular interaction with the catalytic domain ^76^. Upon physical association with permissive membranes, this M domain dissociates, thereby enhancing catalytic efficiency. Biochemical and structural studies of CCT membrane translocation have shown that CCT activation is triggered by biophysical conditions that reduce membrane PC content ^64,67,77^. Thus, CCTs act as highly sensitive molecular sensors of lipid packing defects, signaling the need for immediate PC biosynthesis to enable membrane expansion or to prevent transition of the NE into a porous state ^67,68^. Within this framework, active Rho1 may modulate local lipid composition, specifically the PA/DAG balance, or alter local membrane curvature to create a microenvironment favorable for Pcy1 recruitment and activation under conditions of high demand for PC and GPLs ^16,62^. Alternatively, Rho1 activity may directly influence Pcy1 partitioning between soluble and membrane-associated pools through post-translational modification or via the regulation of nuclear transport machinery ^78^.

Our results also indicate that phospholipid availability differentially affects organelles along the secretory pathway ^65,77^. In the *rgf3*ΔN2 mutant, the KP defect, manifested by reduced PC levels, does not globally disrupt membrane integrity but selectively impacts the NE and the cortical endoplasmic reticulum (ER). This specificity likely reflects intrinsic differences in the biophysical properties of distinct organellar membranes ^79^. The ER, enriched in unsaturated phospholipids and low in sterols, exhibits loose lipid packing that may render it particularly sensitive to phospholipid depletion. Conversely, the plasma membrane (PM), rich in saturated lipids and sterols, displays tight lipid packing and may therefore be more resilient to moderate reductions in PC content ^65,77^.

Like Pct1 in budding yeast, fission yeast Pcy1 localizes to the NE and nucleoplasm (Figure 6F). This spatial confinement could minimize interference from other organelles whose lipid environments may tolerate the unstructured domain of Pcy1 but are not necessarily associated with the high PC demand required for mitotic NE growth ^64^. During mitosis, dividing cells must synthesize sufficient membrane components to increase surface area while strictly preserving proper lipid composition. CCTs contribute to this balance by sensing local PC levels and converting these biophysical signals into changes in enzyme activity and CDP–choline synthesis.

Our data indicate that Rgf3-mediated Rho1 activation coordinates cell-wall septum ingression with mitotic exit by modulating GLP biosynthetic precursors through the Pcy1 “sensor” of the KP. Finally, although mitosis in higher eukaryotes is open rather than closed, their nuclear envelope also undergoes expansion during interphase, immediately following NE reassembly ^7,80^. Because Rho GTPase–regulated processes and the Kennedy pathway of GLP biosynthesis are evolutionarily conserved, studying NE dynamics in yeast provides a valuable, simplified framework for understanding analogous homeostatic processes in higher eukaryotes, including those tightly linked to human tumorigenesis and cellular aging.

## Acknowledgments

We thank H. Valdivieso, S. Oliferenko, I. Hagan, S. Rincón, J.C. Ribas, P. T. Tran, S. Martin, P. Nurse, K. L. Gould, F. Verde, C. Roncero, S. Moreno and P. Pérez for sharing the strains used in this work. We also thank J. Encinar del Dedo for his very helpful comments regarding the manuscript. We are grateful to J. Pinto (bioinformatics facility at the Institute of Biology nd Functional Genomics) for ImageJ macro used for fluorescence quantification. R. Celador and V. Tajadura received a four-year PhD fellowship [University Teacher Training program (FPU)] from the Spanish Ministry of Science and Innovation. P.G. is the recipient of a two-year postdoctoral fellowship (Program II) from the University of Salamanca. T. Edreira received funding through a contract obtained from the Regional Government of Castile and León and co-financed by the European Social Fund.

Funding sources: Spanish Ministry of Economy, Commerce and Business (PID2020-115111GB-I00), the Regional Government of Castile and León [SA116G19], and the NIH (5R35GM149248). Y. Sanchez would also like to thank the institutional support given to the IBFG by the Regional Government of Castile and León through the program “Escalera de Excelencia,” cofinanced by the FEDER Operative Program of Castile y León 2014-2020.

## Author contributions

Conceptualization: R.C. and Y.S.; Methodology, R.C., and P.G.; Experimental research: R.C., V.T., T.E and M.C.; Original manuscript: Y.S.; Writing revision and Editing of the manuscript: R. C., P.G., J. M. and Y.S.; Funding acquisition: Y.S.; Supervision: Y.S and J. M.

## Materials and methods

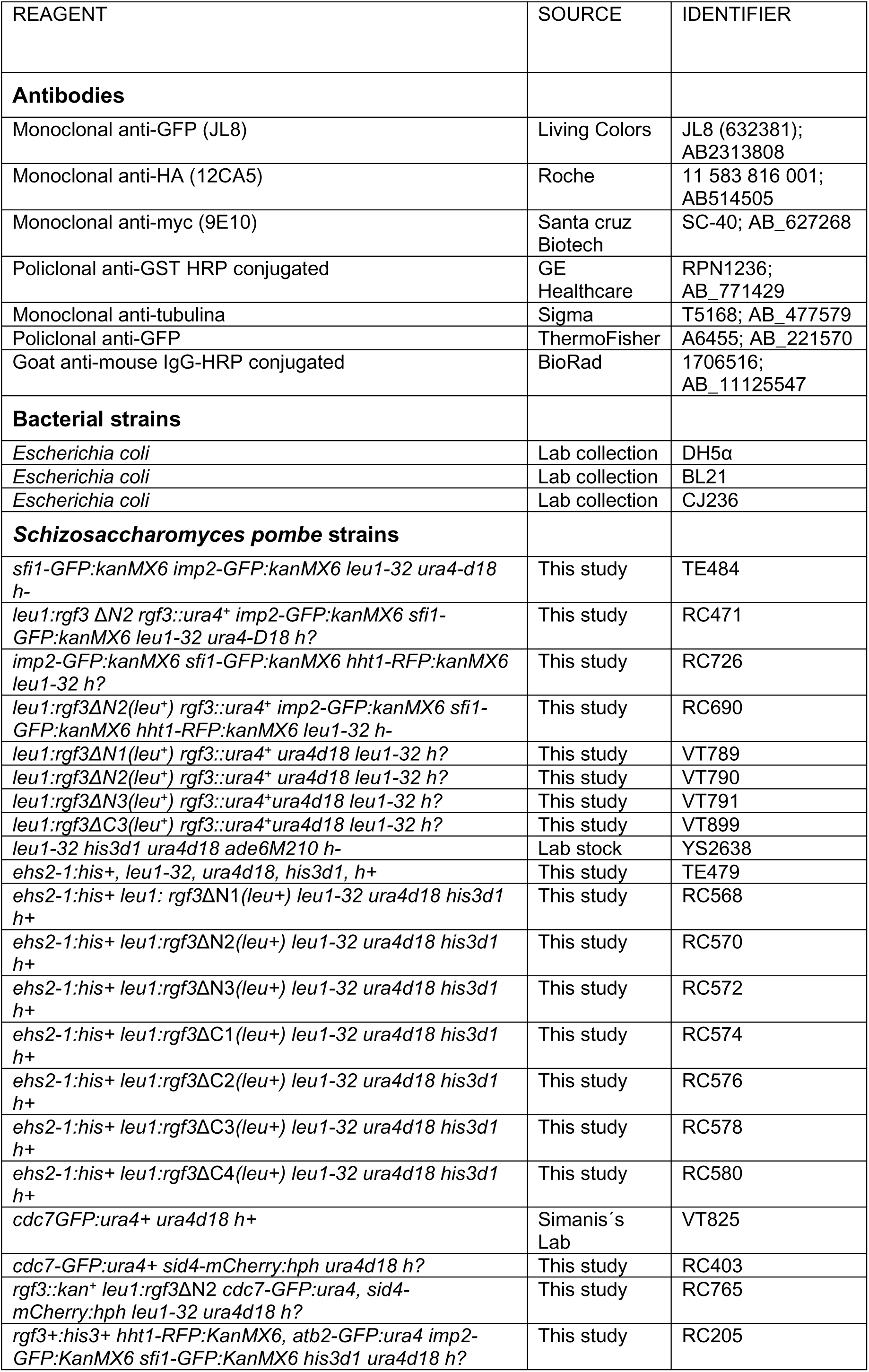
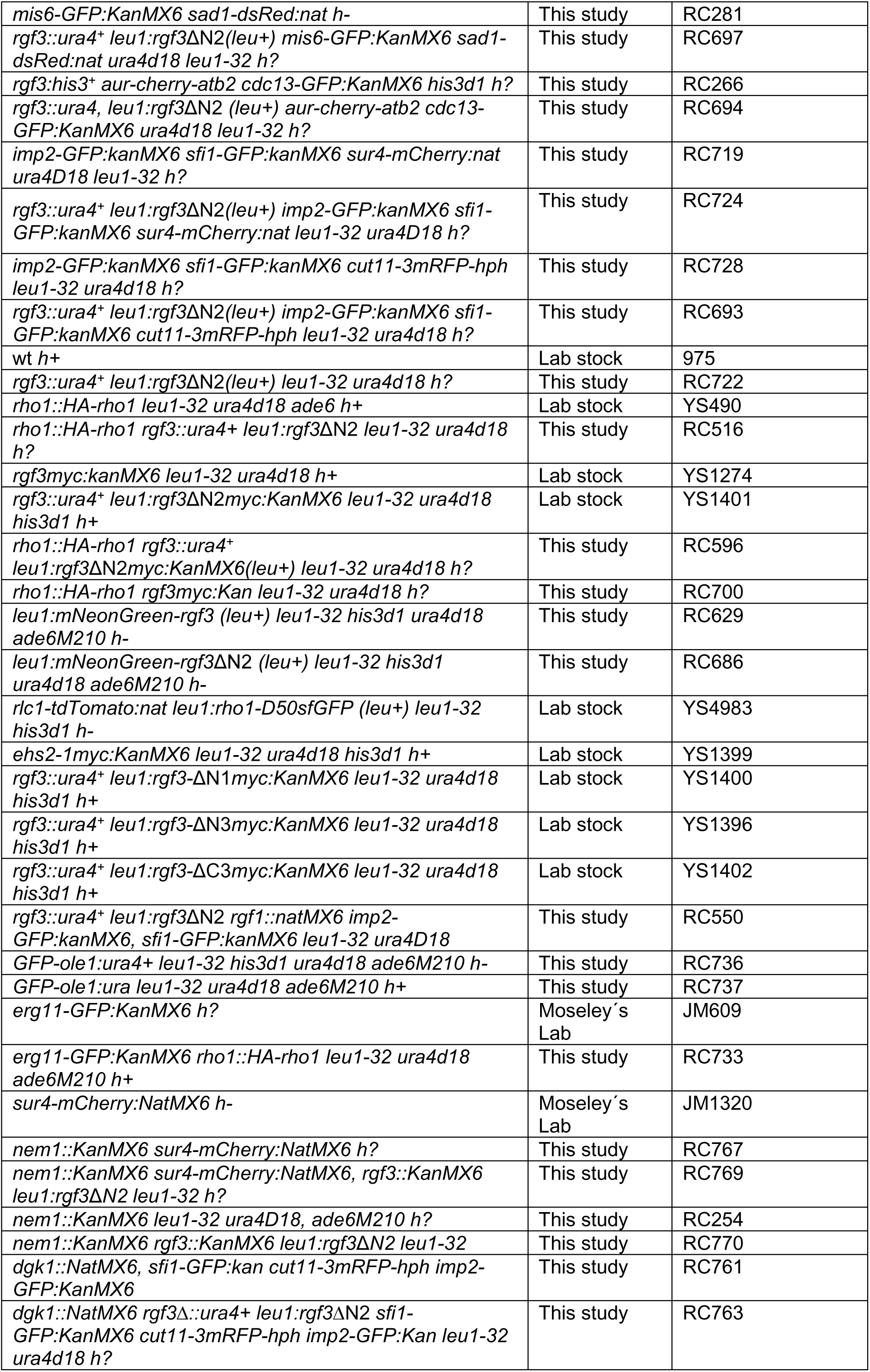
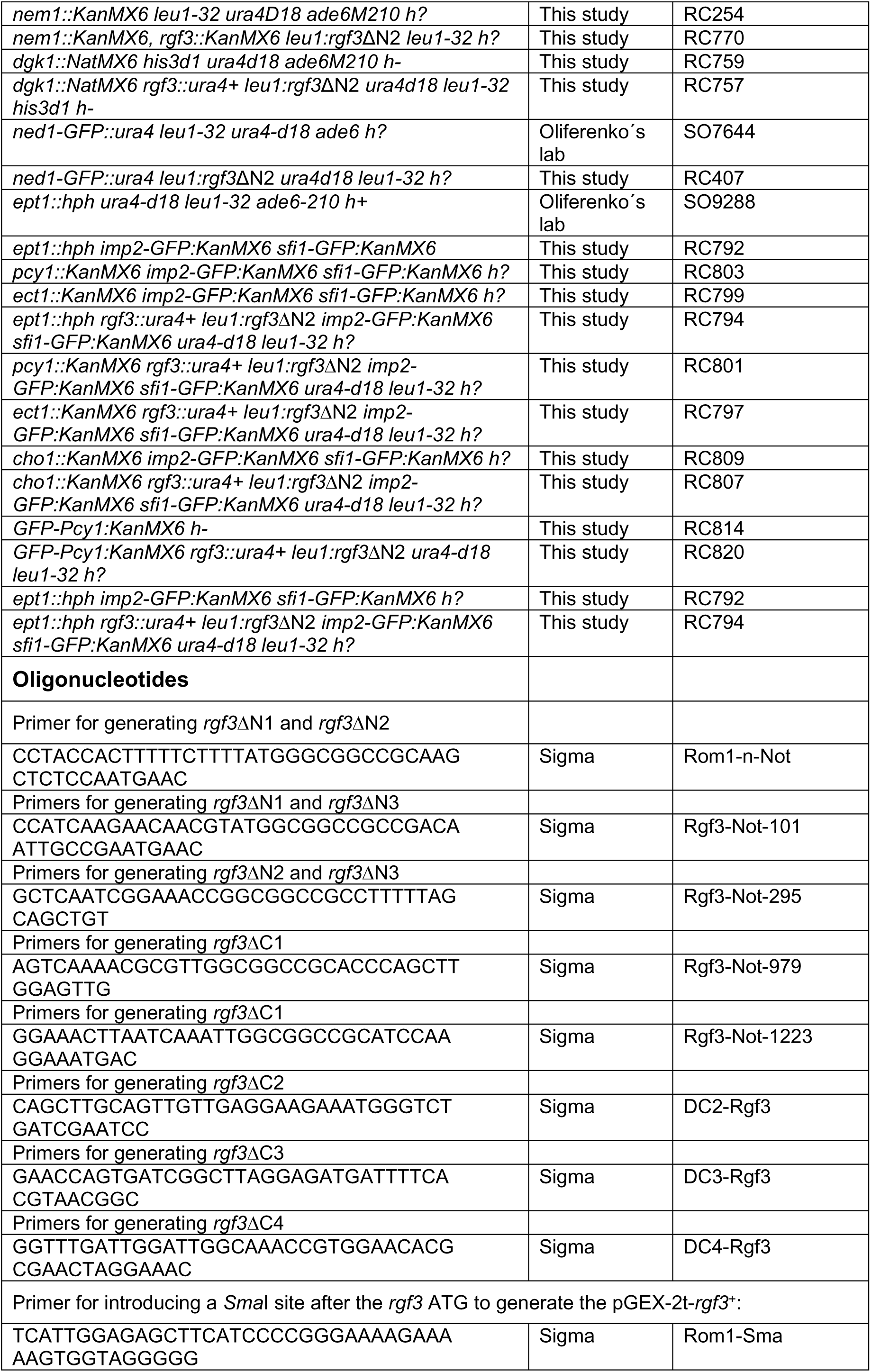
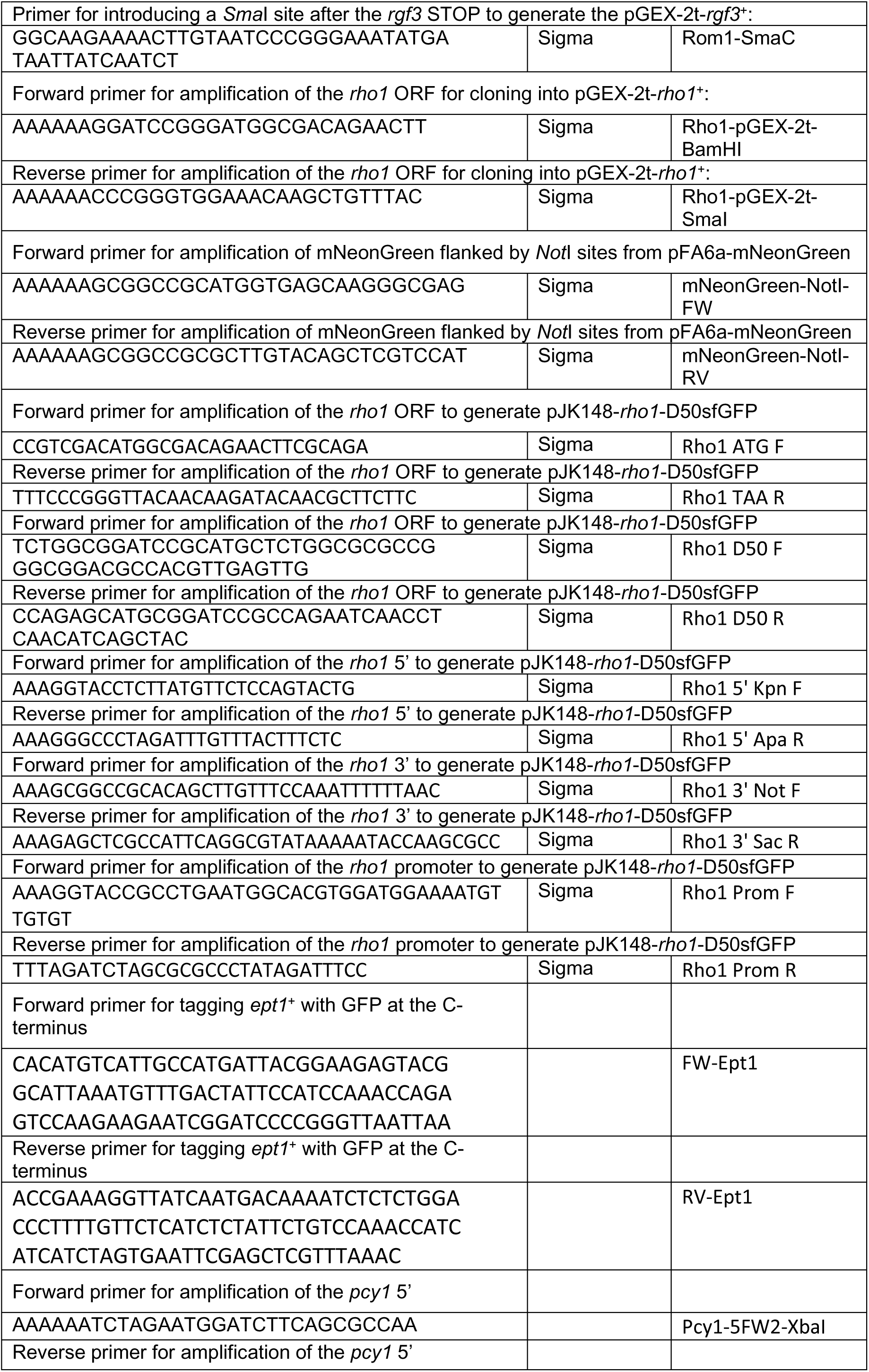
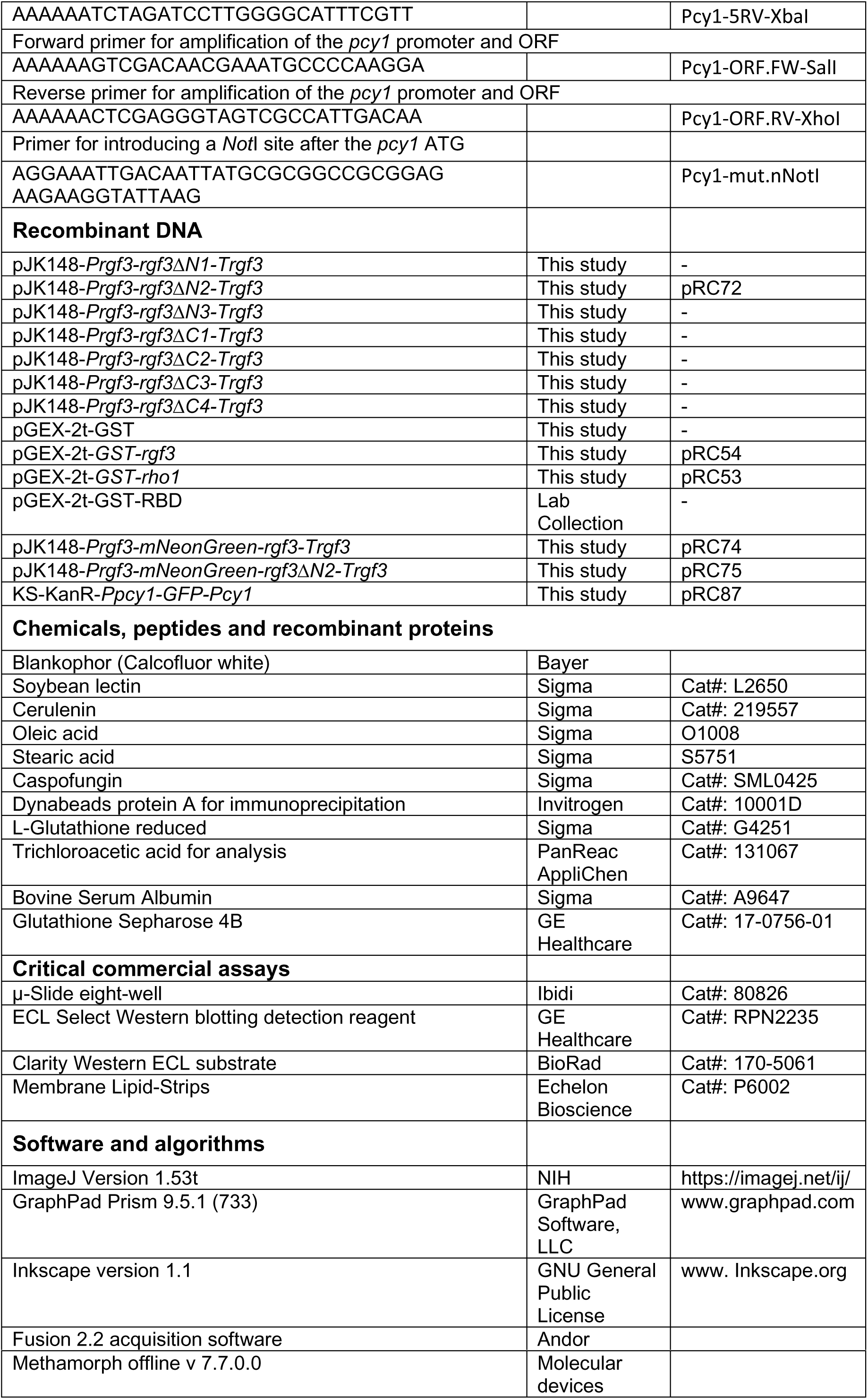

## Resource availability

### Lead contact

Further information and requests for resources and reagents should be directed to and will be fulfilled by the lead contact, Professor Yolanda Sánchez.

### Materials availability

All plasmids/strains generated in this study will be available on request from the lead contact, but may require a completed signed Materials Transfer Agreement.

### Data and code availability

The data reported in this paper will be shared by the lead contact upon request. This paper does not report the original code. Any additional information required to reanalyze the data reported in this paper is available from the lead contact upon request.

### Strains and culture conditions

*S. pombe* strains were streaked onto plates containing complete yeast growth medium (YES) or selective medium (EMM) supplemented with the appropriate requirements and incubated at 28°C until colony formation ^81^. For each biological replicate, a single colony was used to inoculate 5 mL of the respective liquid media. Cultures were incubated at 28°C overnight with shaking (200 rpm). Each overnight culture was subsequently used as a seed culture to inoculate fresh media. Fresh cultures were grown at 28°C at 200 rpm and to an OD_600_ of 0.5-0.6 at the time of harvesting the cells. Geneticin, hygromycin, and nourseothricin were used at 120 μg/ml, 400 µg/ml, and 50 μg/ml, respectively. Cerulenin (stock at 5 mg/ml in DMSO) was used at 5 μM in liquid culture and 0.5 μg/ml in the drop assays. Cerulenin, caspofungin, and other drugs were added to the medium after autoclaving. Crosses were performed by mixing appropriate strains directly on MEA plates. Recombinant strains were obtained by tetrad analysis or the “random spore” method. For fatty acid supplementation, oleic acid or stearic acid was added to a concentration of 1 mM to MM from a 12.75 mM stock in 12% (w/v) fatty acid-free BSA ^82^. *BSA conjugation of free fatty acids*. Fatty acid-free BSA (24% w/v) was made by adding 12 g of fatty acid-free bovine serum albumin to 35 ml of 150 mM NaCl in six 2-gram doses over 5 hours. The pH was adjusted to 7.4 with 5 M NaOH, and the volume was brought up to 50 ml with 150 mM NaCl. This solution was diluted 1:1 with 150 mM NaCl to produce 12% fatty acid-free BSA before use as an experimental control. Fatty acid conjugation to BSA was performed by dissolving 319 µmol of oleic acid or stearic acid in 2 ml of ethanol. In total, 100 µl of 5 M NaOH was added to precipitate the sodium salt of the fatty acid and the ethanol was then evaporated in a SpeedVac. Subsequently, 10 ml of 150 mM NaCl were added to the dried fatty acid, and the solution was heated until the fatty acid dissolved. The solution was then stirred and cooled, avoiding fatty acid precipitation, at which point 12.5 ml of ice-cold fatty acid-free BSA (24% w/v) was added. The solution was then stirred for ten minutes, and the final volume was adjusted to 25ml with 150 mM NaCl.

### Plasmid and DNA techniques

Plasmids *rgf3*ΔN1*, rgf3*ΔN2*, rgf3*ΔN3*, rgf3*ΔC1*, rgf3*ΔC2*, rgf3*ΔC3*, and rgf3Δ*C4 originate from the pYSJK5 plasmid backbone. pYSJK5 is pJK148-*rgf3*^+^ (integrative vector pIJ148) with its promoter and terminator sequences. For each construct, we introduced two *Not*I sites flanking the region to be deleted by site-directed mutagenesis. Plasmids were linearized and inserted into the *leu1* locus of a diploid strain (*rgf3*^+^/*rgf3*::*ura4*). After sporulation, the colonies able to grow in MM-ura and MM-leu, which carried the deletion of Rgf3 and the corresponding N-terminus deletion mutant, were selected. Clones from *rgf3*ΔN1 (deletion of aminoacids 1-101), *rgf3*ΔN2 (1-295), *rgf3*ΔN3 (101-295), and *rgf3*ΔC3 (1071-1090, 20 aminoacids deleted within the central region of the CNH domain) were established as haploids. Labelling the full–length *rgf3*^+^ and mutants with 13myc at the C-terminus was done using long oligonucleotides as indicated in ^83^. To detect Rho1-GTP levels, we used the pGEX-C21RBD plasmid (rhotekin-binding domain) kindly provided by P. Pérez (IBFG, Salamanca, Spain). The proteins were expressed in *Escherichia coli* using pGEX-2T (Sigma-Aldrich). For the lipid strip assays, pGEX-*rgf3^+^* was made by inserting the ORF of *rgf3* (obtained by introducing a *Sma*I site before the ATG and one *Sma*I site right after the END by site-directed mutagenesis in pYSJK5) at the *Sma*I site of pGEX-2T. pGEX-*rho1^+^* was obtained by inserting the ORF of *rho1^+^* (without introns and flanked by *Sma*I-*BamH*I sites), in frame, into the same restriction sites of pGEX-2T. The mNG-Rgf3 strain was obtained by integrating the pRC74 (pJK148-mNeonGreen-*rgf3*^+^) into the *leu1* locus. pRC74 was derived from pRC3 (pJK148-GFP-*rgf3*+) by replacing the GFP with a *Not*I-flanked mNeonGreen sequence amplified from pFA6a-mNeonGreen-KanMX6. pRC3 was generated by inserting a GFP into a *Not*I site introduced downstream of the *rgf3* start codon by site-directed mutagénesis. The strain expressing the truncated fusion mNG-*rgf3*-ΔN2 was obtained by integrating plasmid pRC75 into the *leu1* locus. pRC75 is derived from pRC72 (pJK148-*rgf3*-ΔN2), in which the mNG sequence was inserted at a specific *Not*I site created upon removal of the N-terminal region, linking the start codon directly to residue 295. sfGFP-*rho1* strain was obtained by integrating plasmid pTE67 (pJK148-*rho1*-D50sfGFP) into the *leu1* locus. pTE67 contains the *rho1* cDNA sequence driven by its native promoter and terminator. The sfGFP tag was inserted internally after residue D50, flanked by the linker sequence SGGSACSGAPG. Labelling *ept1*^+^ with GFP at the C-terminus was done using long oligonucleotides as indicated in ^83^. The GFP-Pcy1 strain was generated by integrating the pRC87 plasmid at the *Pcy1* locus. The 5’ region of Pcy1 was obtained by PCR and cloned into a KS-*Kan*^R^ plasmid within the *Xba*I site to yield pRC84. A second PCR fragment, containing the promoter and part of the Pcy1 ORF, was then inserted between the *Sal*I and *Xho*I sites of pRC84 to obtain pRC85. pRC85 was modified by site-directed mutagenesis to introduce a *Not*I site after the Pcy1 ATG, which was used to clone the GFP flanked by NotI sites, yielding pRC87. A second strategy allowed us to overexpress Pcy1 fused to GFP at its N-terminus by replacing its native promoter with the *nmt41x* promoter. For this purpose, long oligonucleotides were used as described in ^83^.

### Survival assays

To test sensitivity to different drugs, cells were grown in YES plates for 2 days (to the stationary phase). Cells were resuspended in sterile water and spotted as serial dilutions, 8 × 10^4^ cells in the left row, and then 4 × 10^4^, 2 × 10^4^, 2 × 10^3^, 2 × 10^2^ and 2 × 10^1^ in each subsequent spot, onto YES plates or YES supplemented with different drugs. The cerulenin sensitivity assay was performed on minimal medium (MM) plates supplemented with all amino acids. All survival assays were carried out in triplicate and, unless otherwise stated, the plates were incubated for 3 days for the YES assays and 5 days for the MM assays at 28°C.

### Protein extraction and Western blot analysis

The cells of *S. pombe* cultures (25 mL) at an OD_600_ of 0.5 were harvested by centrifugation at 4°C at 1690 g for 3 min, washed once with 1 ml of cold TCA 20%, suspended in 100 μl of 20% TCA and frozen in liquid N_2_. Acid-washed glass beads were added, and the cell homogenates were prepared in a ‘Fast Prep’ FP120 device (Savant; Bio101). Four hundred microliters of cold 5% TCA were added to each tube, which was vortexed to wash the beads. Cell extracts were transferred to a clean tube and centrifuged for 10 minutes at 1690 g at 4°C. The pellets were resuspended in a 2% sodium dodecyl sulfate (SDS)/0.3 M Tris base. The samples were boiled and clarified by centrifugation at 16000 g for 1 minute. Protein concentration was determined using the Bradford protein assay (Bio-Rad), and the samples were mixed with Laemmli sample buffer (50 mM Tris-HCl, pH 6.8; 1% SDS; 143 mM β-mercaptoethanol; 10% glycerol). Samples were subjected to polyacrylamide gel electrophoresis (5.5% gels for Rgf3 and 12% for Rho1), transferred to PVDF membranes, and incubated in blocking buffer (5% Nestlé non-fat dried milk in TBST: 0.25% Tris, pH 6.8; 0.9% NaCl; 0.3% Tween 20) for 1 hour. Primary antibodies were anti-myc (9E10, Santa Cruz Biotechnology), anti-GFP (JL8, Living Colors; 1∶3000), anti-HA (12CA5 mAb, 1:5000), anti-GST (RPN1236V, Citiva, 1:5000), and anti-α tubulin (clone B-5-1-2; 1∶10000). The secondary antibody was horseradish peroxidase-conjugated anti-mouse (1∶10000). Chemiluminescent signals were detected with a Vilber Fusion FX using the Clarity Western ECL substrate (BioRad).

### Protein immunoprecipitation

For the immunoprecipitation assays, cells from 100 ml cultures at an OD_600_ of 0.6 were harvested and washed in 2 ml of STOP buffer (150 mM NaCl, 50 mM NaF, 10 mM EDTA, and 1 mM NaN_3_). The pellets were re-suspended in lysis buffer (pH 7.5, 50 mM HEPES, 100 mM NaCl, 1mM EDTA, 20 mM β-glycerophosphate, 50 mM NaF, 1% NP-40, 1 mM phenylmethylsulfonyl fluoride, 1 μg/ml leupeptin, 1 μg/ml aprotinin) and lysed by bead disruption. The extracts were cleared by centrifugation at 10000 rpm for 10 min and 3000 μg of total protein were used in each immunoprecipitation reaction in a final volume of 1 ml. The Rgf3 and ΔN mutants were immunoprecipitated from lysates using an anti-c-myc polyclonal antibody (C3956; Sigma) and protein A Dynabeads (10001D, Invitrogen). The beads were washed three times with lysis buffer and suspended in 25 µL of Laemmli buffer.

### *In vitro* production of recombinant proteins and the pull-down assay

For the pull-down and lipid strip assays, first GST-tagged Rgf3, Rho1 or C21RBD (rhotekin-binding domain to detect Rho1-GTP levels) were produced from *E. coli*. The fusion proteins were induced by the addition of 0.5 mM IPTG at 18°C overnight (or 3 hours at 28°C for C21RBD). Cells were sonicated and proteins were immobilized on glutathione-Sepharose 4B beads (GE-Healthcare). After one hour of incubation, the beads were washed three times, and the bound proteins were analyzed by SDS-PAGE and stained with Coomassie brilliant blue. The amount of GTP-bound Rho1p was determined using a pull-down assay as described previously ^84^. In brief, extracts from 50-ml cultures of wild-type and *rgf3*ΔN2 strains carrying HA-Rho1 expressed from its promoter were re-suspended in 500 μl of lysis buffer (50 mM Tris, pH 7.5, 20 mM NaCl, 0.5% NP-40, 10% glycerol, 0.1 mM dithiothreitol and 2 mM Cl_2_Mg, containing 100 μM PMSF, leupeptin, and aprotinin). After beating, cell extracts (2 mg of total protein) were incubated with 10 μg of GST–C21RBD coupled to GS beads for 2 hours, washed four times, and bound proteins were blotted against 1:5000-diluted 12CA5 mAb to detect HA-Rho1. Total levels of HA-Rho1p were monitored in whole-cell extracts (50 μg of total protein) by Western blot analysis.

### Lipid strip overlay

Overlays were performed as previously described {Fernandez-Golbano, 2014 #6412} using lipid strips (p-6002, Echelon). Strips were blocked with 3% fatty acid-free BSA (Sigma) in TBS-T buffer (10 mM Tris pH 8.0, 150 mM NaCl, 0.1% Tween-20; TBST-BSA) at room temperature for 1 hour and incubated further for 2 hours together with 1 μg/mL GST, GST-Rgf3 or GST-Rho1 in TBST-BSA buffer. The strips were then washed 3 times with 5 ml of TBST-BSA and incubated with Anti-GST HRP conjugate antibody (Cytiva) in TBST-BSA. Bound protein was detected using an enhanced chemiluminescence detection kit (BioRad). GST, GST-Rgf3, and GST-Rho1 proteins were expressed in *E. coli* and purified using glutathione-Sepharose beads (GE-Healthcare) according to the manufacturer’s instructions and as described in the previous section. Once attached to the beads, the proteins were washed and eluted with 200 μL of elution buffer (100 mM Tris-HCl, pH 8.0, 120 mM NaCl) with 20 mM of L-Glutathione reduced (Sigma) freshly added for 30 min at 4°C. Aliquots were frozen in liquid air with 15% glycerol and stored at −80°C until used.

### Lipidomics

Cells of the WT strain and the *rgf3*ΔN2 mutant (OD600 ∼ 1.0) grown in MM were collected by centrifugation (at 1690 g for 3 min at 4°C) and washed three times with ultrapure water (Milli-Q IQ7005; Millipore, Bedford, MA, USA). The cell pellets were then suspended in 300 µL of ultrapure water. Glass beads (1.0 g; acid-washed, 0.4 mm diameter) were added and the mixtures were shaken 3 times for 40 seconds at 6.5 m/s each time in a FastPrep-24 5G. Finally, the homogeneous broken cell suspensions were collected, diluted with ultrapure water (∼ 20 OD_600_ units/ml) and subjected to a lipidomics analysis. Sample extracts, fortified with internal standards, were prepared by mixing 100 μL of cell lysates with 0.75 mL of methanol and chloroform (2:1 v/v) containing 0.01% BHT. The samples were sonicated in a bath sonicator until evenly dispersed, then incubated overnight at 48°C in a water bath. After cooling, 75 μL of 1M KOH in methanol was added, and the samples were sonicated and incubated for 2 hours at 37°C. Solvents were removed using a SpeedVac concentrator, and the samples were redissolved in methanol and transferred to a new tube. Solvents were removed again with the SpeedVac concentrator. The dried samples were redissolved in methanol, centrifuged, and the supernatants were transferred to a glass vial for HPLC analysis.

Liquid chromatography-mass spectrometry was performed using a Waters Acquity Premier UPLC system coupled to a SELECT SERIES Cyclic IMS, which combines a Q-Tof (Quadrupole-Time of Flight) mass spectrometer with cIM (Cyclic Ion Mobility) (Waters, Milford, MA), operated in positive electrospray ionization mode. Full scan spectra from 50 to 1200 Da were acquired, and individual spectra were summed to generate data points every 0.5 s. Mass accuracy and reproducibility were maintained using an independent reference spray via the LockSpray interference. The analytical column was a 100 mm × 2.1 mm i.d., 1.7 μm C8 Acquity UPLC BEH column (Waters). The two mobile phases were: phase A: water with 2 mM ammonium formate; phase B: methanol with 1 mM ammonium formate, both containing 0.2% formic acid. A linear gradient was programmed as follows: 0.0 min: 80% B; 3 min: 90% B; 6 min: 90% B; 15 min: 99% B; 18 min: 99% B; 20 min: 80% B; 22 min: 80% B. The flow rate was 0.3 mL/min, and the column was held at 30°C. Quantification was performed using the extracted ion chromatogram of each compound with 50 mDa windows. The linear dynamic range was determined by injecting standard mixtures. Positive identification of compounds was based on accurate mass measurement with an error of <5 ppm and their LC retention time, compared to that of a standard.

### Microscopy and image analysis

Fluorescence images were acquired on a Nikon Eclipse Ti2 Andor Dragonfly 200 Spinning-disk confocal microscope equipped with a sCMOS Sona 4.2B-11 camera (Andor) and controlled with Fusion 2.2 acquisition software (LLC). Approximately 1 ml of early log phase cells was collected by applying a short spin and dissolved in 5 µL of water. Then, 10 slices at 0.5 µm were taken, corrected by 3D Deconvolution (conservative ratio, 10 iterations and medium noise filtering) with Fusion software (Oxford Instruments), and processed with Fiji distribution of Image {Schindelin, 2012 #5800}.

SRRF super-resolution images were acquired on a Nikon Eclipse Ti2 microscope coupled to an Andor Dragonfly 200 spinning disk confocal system, using a 100× objective and an additional 1.5× magnification lens. A total of 25 images were collected at the midplane for each strain analyzed.

To analyze protein or organelle dynamics, time-lapse imaging was carried out separately on two different microscopes: an Olympus IX81 spinning disk microscope (Olympus) equipped with a PlanApo 100x/1.40 oil immersion objective, a confocal CSUX1-A1 module (Yokogawa), and an Evolve (Photometrics) camera controlled by MetaMorph software (Molecular Devices) and a Nikon Eclipse Ti2 Andor Dragonfly 200 Spinning-disk confocal microscope equipped with a sCMOS Sona 4.2B-11 camera (Andor) and controlled with Fusion 2.2 acquisition software; LLC). One mL of cell cultures at mid-log phase was collected by centrifugation (at 3500 g for 1 min), resuspended in 0.3 ml of YES containing the respective drugs (BP and cerulenin), and placed in the wells of a µ-Slide eight-well plate (Ibidi). Each of the wells was previously coated with 5 µL of 1 mg/mL soybean lectin (Sigma-Aldrich). The sum projection of a 10-plane stack was used. The cells were allowed to adhere to the bottom of each well for 2.5 min before removing the medium. Then the cells were washed three times with medium and suspended in 0.3 ml of the same medium. Time-lapse experiments were performed at 32°C. For imaging the temperature-sensitive strains at higher temperatures, the cells were shifted from 25°C to the restrictive temperature and incubated for 3 hours under these conditions before imaging. Then, 10 slices at 0.5 µm were taken every 3 min in up to four regions in each of the four different wells.

In most of the time-lapse series, we used the SPB separation as time zero of the cellular clock. We define cytokinesis completion time as the time between SPB separation and complete closure of the AR. The percentage of cells with problems segregating was calculated considering only dividing cells. Cells whose nuclei were cut by the biosynthetic machinery, cells in which both nuclei ended up on the same side of the daughter cell or those in which the daughter nuclei had asymmetric sizes were considered missegregating cells. The length of the mitotic spindle was calculated indirectly as the distance that separates the two SPBs along the time-lapse series; images were taken every 1.5 minutes for 3 hours. The maximum projection of a 10-plane stack was used to measure the area of the nucleus from the time when two SPBs could be distinguished until the nuclei reached the cell poles. A line was drawn following the circumference of the nucleus in each frame, and the internal area of the figure was calculated using Fiji (Fiji is Just Image J). The ratio of Pcy1 fluorescence between the nuclear envelope and the nucleoplasm was quantified by measuring the mean fluorescence intensity in 0.065 µm² rectangular regions selected within each compartment. The final ratio was calculated as the fluorescence of the envelope divided by that of the nucleoplasm.

## Quantification and statistical analysis

Statistical analyses and graphs were generated using GraphPad Prism Software version 9.5.1. To compare two conditions, a two-tailed unpaired Student’s t-test was applied to determine statistical significance (as detailed in the figure legends), and values of P < 0.05 were considered as being significant. The graphs show the mean ± standard deviation of the indicated data. Asterisks represent the following: *P < 0.05 **P < 0.01; ***P < 0.001 ****P < 0.0001.

---

**Figure S1.**
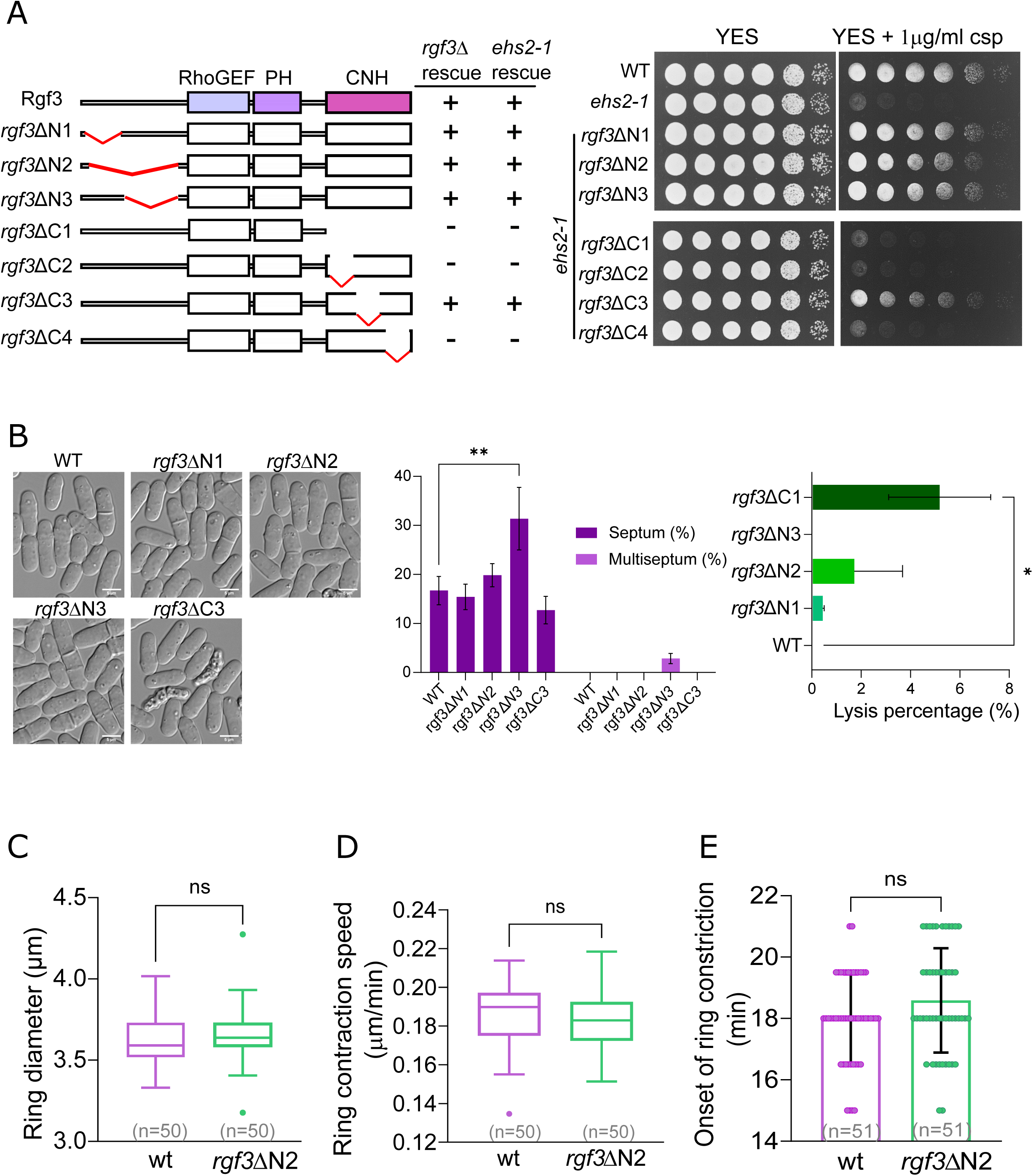
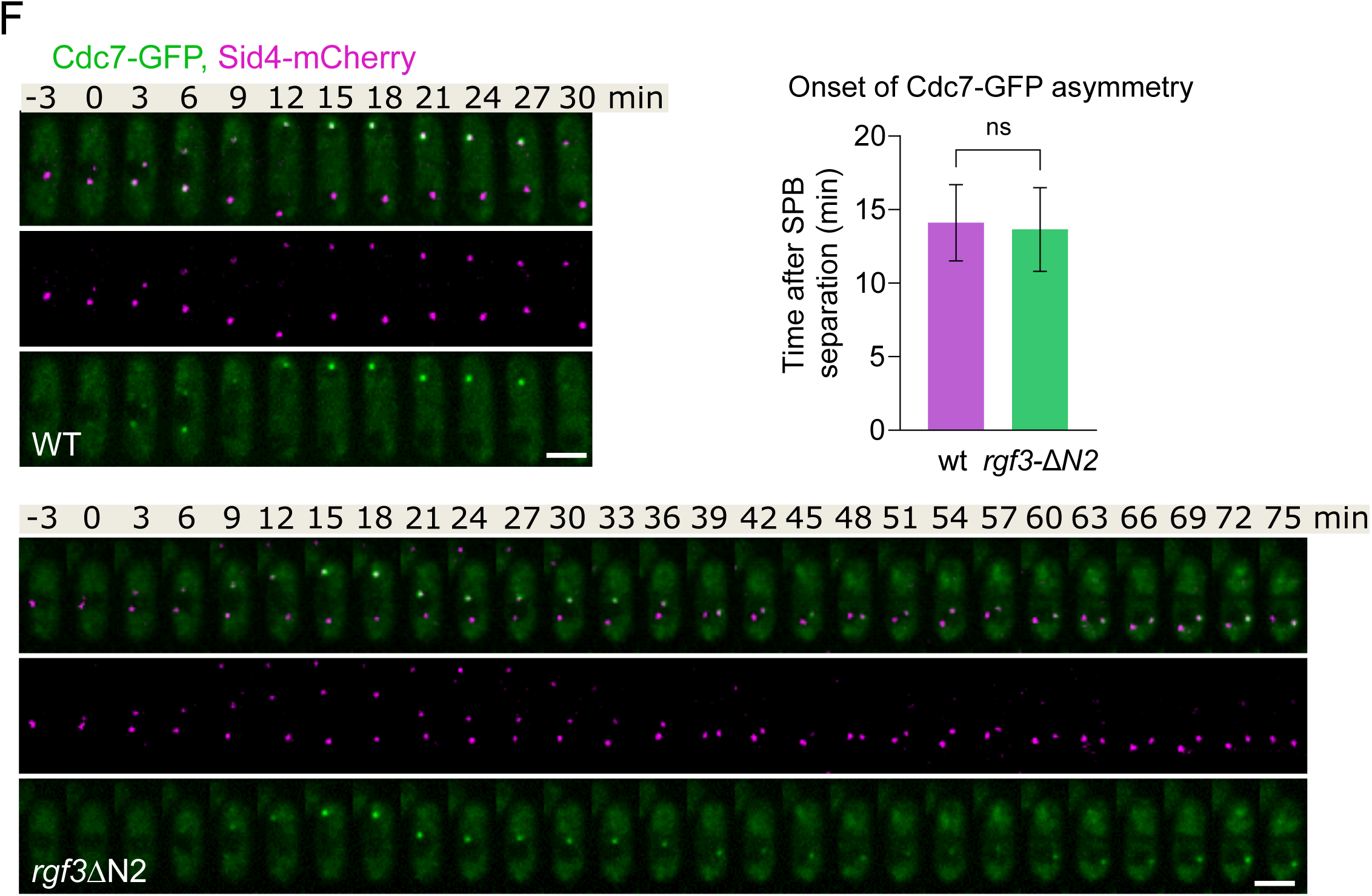
The N-terminal domain of Rgf3 is dispensable for cell viability but required for proper mitotic progression. **(A)** The N-terminal domain of Rgf3 is non-essential for cell viability. Full-length Rgf3 and various deletion fragments were integrated at the *leu1* locus in a diploid (*rgf3*Δ/*rgf3*^+^) and a haploid *ehs2-1* background. Diploids were sporulated, and deletion constructs capable of rescuing *rgf3*Δ lethality are indicated by (+). For the *ehs2-1* rescue assay, the ability of the mutant fragments to suppress Caspofungin (Csp) sensitivity is indicated by (+) and shown in the drop assay on the right. The indicated strains (grown in YES) were adjusted to an initial O.D._600_ of 2.7. A series of three two-fold serial dilutions followed by two ten-fold serial dilutions were spotted onto rich YES plates with or without caspofungin (1µg/ml). Colony formation was analyzed after 2–3 days of incubation at 28°C. **(B)** (Left panel) Live-cell differential interference contrast (DIC) micrographs of wild-type (WT), *rgf3*ΔN1, *rgf3*ΔN2, *rgf3*ΔN3, and *rgf3*ΔC3 cells grown to log phase in YES medium. Scale bars, 5 µm. (Middle panel) Percentage of septated cells (one septum or ≥2 septa) for the indicated strains grown in YES at 28°C. To visualize septa, cells were stained with Blankophor (BP, 1 µg/ml) immediately prior to imaging. Data represent the mean ± SD from three independent experiments (n > 200 cells analyzed per strain). (Right panel) Quantitation of cell lysis percentages. Log-phase cells were scored under light microscopy to evaluate the proportion of lysed cells. Data represent the mean ± SD from three independent experiments (n = 100 cells per strain). **(C and D)** Ring dynamics quantification from individual cells shown in Figure 1F (n = 50 cells per strain), with time 0 min defined as SPB separation. (C) Mean ring diameter prior to the onset of contraction, and (D) ring contraction speed (µm/min) in wild-type (WT) and *rgf3*ΔN2 strains. Boxes represent the interquartile range (IQR), whiskers depict Tukey’s range, and the horizontal line inside the box indicates the median value (n.s., non-significant; calculated by an unpaired Student’s t-test). **(E)** Time from SPB separation to the onset of ring constriction measured in WT and *rgf3*ΔN2 cells (n = 51 cells per strain). Bar plots represent the mean ± SD (n.s., non-significant; calculated by an unpaired Student’s t-test). **(F)** Representative time-lapse fluorescence micrographs of wild-type (WT) and *rgf3*-ΔN2 cells expressing Cdc7-GFP (green) and Sid4-mCherry (magenta; SPB marker), captured via spinning-disk confocal microscopy. (Right graph) Quantitation of the time from SPB separation to the onset of Cdc7-GFP asymmetry (n = 20 cells analyzed per strain). Bar plots represent the mean ± SD (n.s., non-significant; calculated by an unpaired Student’s t-test). Scale bars, 5 µm.

**Figure S2.**
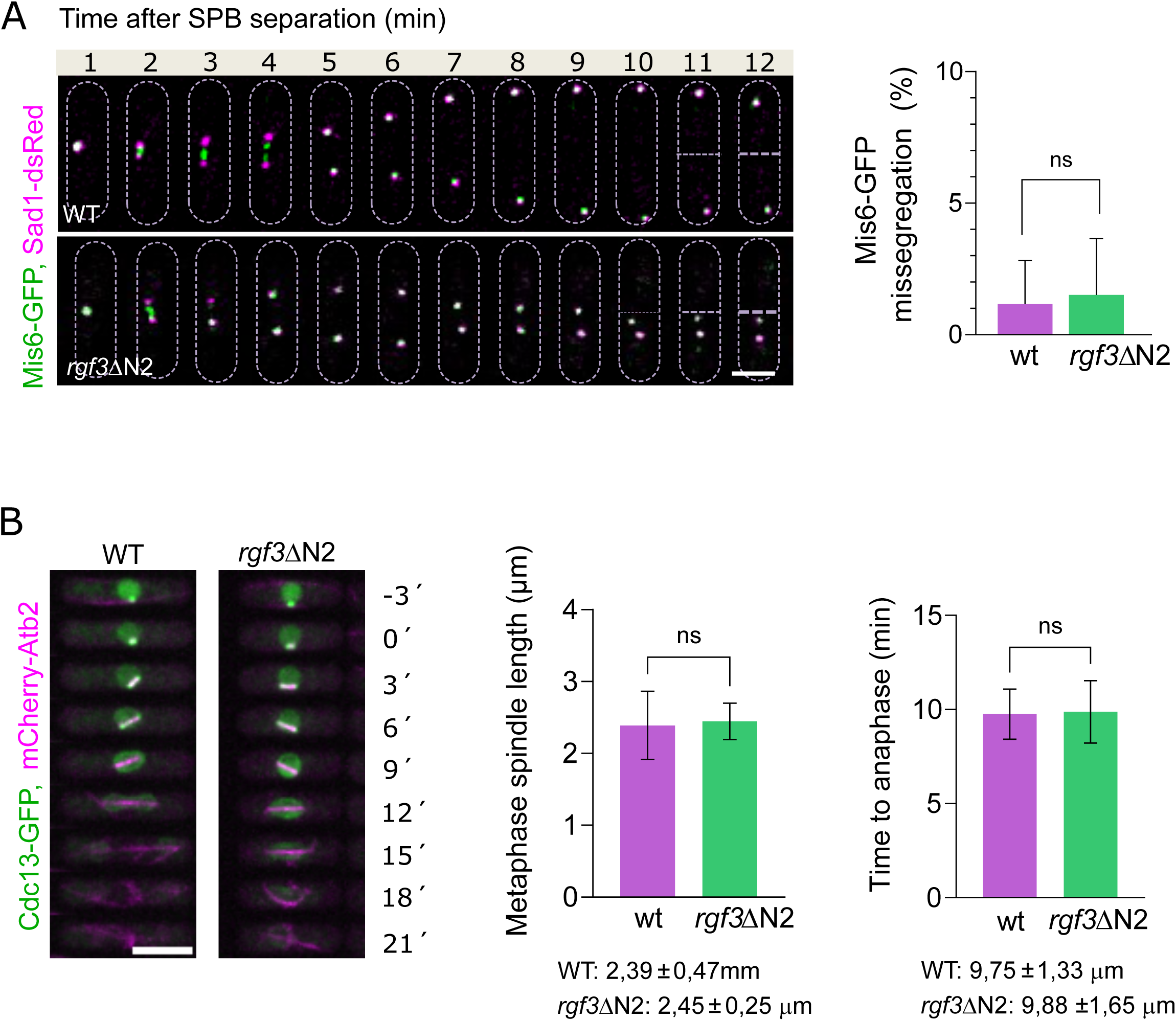
**(A)** (Left) Time-lapse micrographs of representative wild type and *rgf3*-ΔN2 cells expressing Sad1-dsRed (SPB marker) and Mis6-GFP (inner kinetochore marker) during mitosis. Images were captured at 1-minute intervals. Scale bar, 5 µm. (Right) Quantification of Mis6-GFP missegregation (%) in WT and *rgf3*-ΔN2 strains. Data represent the mean ±SD from two independent experiments (>20 cells analyzed per strain). Statistical significance was determined by an unpaired *t*-test (n.s., non-significant). **(B)** (Left) Time-lapse micrographs of representative wild-type (WT) and *rgf3*-ΔN2 cells expressing mCherry-Atb2 (α-tubulin) and Cdc13-GFP (cyclin B) from the onset of mitosis to spindle breakdown. Images were captured at 3-min intervals. (Middle) Metaphase spindle length (µm) at the transition to anaphase (defined by Cdc13-GFP degradation) in WT (purple, n = 24) and *rgf3*-ΔN2 cells (green, n = 24). (Right) Time from mitosis onset to anaphase transition (min) in WT (purple, n = 24) and *rgf3*-ΔN2 cells (green, n = 24). Values below the graphs indicate the mean ± SD for each strain. Data are presented as mean ± SD, and *p*-values were calculated using a Student’s *t*-test (n.s., non-significant). Scale bar, 5 µm.

**Figure S3.**
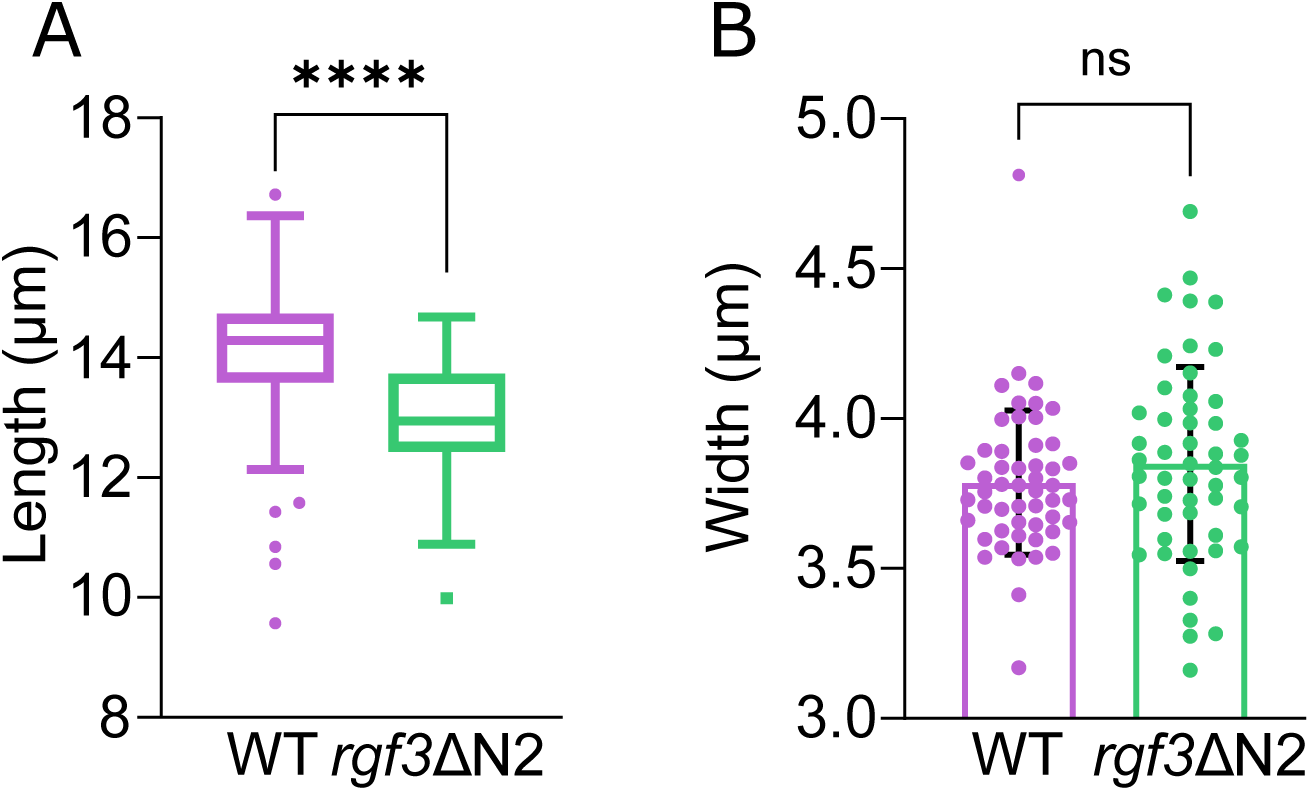
Cell length and width analysis of wild-type and rgf3ΔN2 strains. **(A)** Box-and-whisker plot displaying the cell length (µm) of septated wild-type (WT; purple, n = 51 cells) and *rgf3*ΔN2 mutant (green, n = 51 cells) strains. Boxes define the interquartile range (IQR), whiskers represent Tukey’s range, and the horizontal line inside the box indicates the median value. **(B)** Bar plot with superimposed individual data points showing the cell width (µm) of WT (n = 51 cells) and *rgf3*ΔN2 (n = 51 cells) strains. Bar heights and error bars represent the mean ± SD. Statistical significance between genotypes was determined using an unpaired Student’s t-test (****, p < 0.0001; n.s., non-significant).

**Figure S4.**
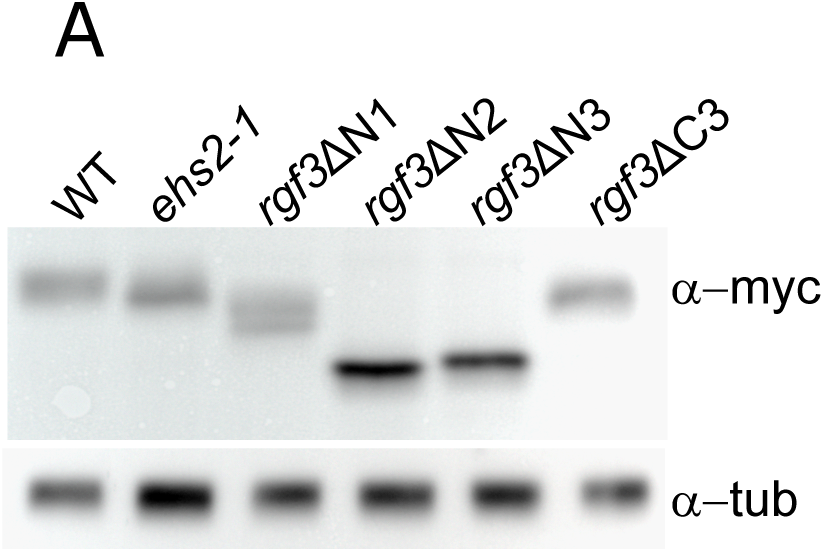
Expression levels and biochemical stability of Rgf3 mutant variants. **(A)** Trichloroacetic acid (TCA) total cell extracts were prepared from asynchronous cultures of wild-type (WT; *rgf3*^+^-myc), *ehs2-1-myc*, *rgf3*ΔN1-myc, *rgf3*ΔN2-myc, *rgf3*ΔN3-myc and *rgf3*ΔC3-myc strains. Protein samples were resolved by SDS-PAGE and subjected to Western blot analysis using an anti-myc antibody (top panel). Immunoblots against α-tubulin (bottom panel) were utilized as a loading control to ensure equal protein concentration across all lanes.

**Figure S5.**
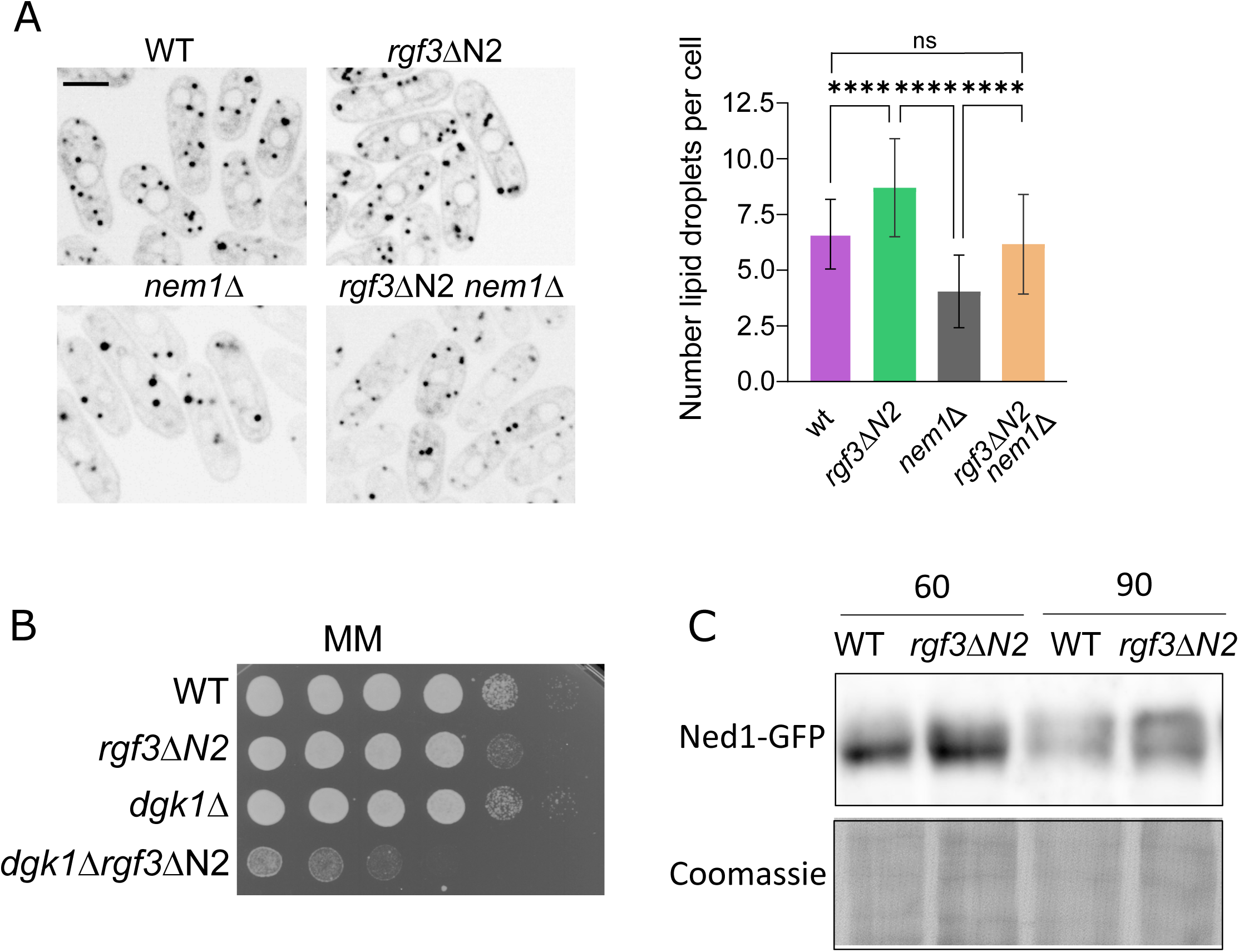
Lipid droplet quantification, genetic interaction assays, and Ned1 phosphorylation status. **(A)** Representative single-plane spinning-disk confocal micrographs of wild-type (WT), *rgf3*-ΔN2, *nem1*Δ and *rgf3*-ΔN2 *nem1*Δ cells showing lipid droplets (LDs) stained with BODIPY 558/568. Scale bar, 5 µm. (Right graph) Quantitation of the number of lipid droplets per cell in the indicated strains. Data represent the averages from three independent experimental replicates (n > 20 cells analyzed per strain). Error bars indicate the SD of the mean. Statistical significance was determined using a one-way ANOVA followed by Tukey’s multiple comparisons test (****, p<0.0001; n.s., non-significant). **(B)** Drop dilution assay showing the synthetic genetic interaction between *rgf3*ΔN2 and *dgk1*Δ. Cells from WT, *rgf3*ΔN2, *dgk1*Δ, and *rgf3*ΔN2 *dgk1*Δ strains were adjusted to an initial O.D._600_ of 2.7. A series of three two-fold serial dilutions followed by two ten-fold serial dilutions were spotted onto minimal medium (MM) plates and incubated for 3–4 days at 28°C. **(C)** Immunoblot analysis of Ned1-GFP phosphorylation profiles. Asynchronous cultures of *ned1-GFP* and *ned1-GFP rgf3*ΔN2 cells were synchronized and blocked in early S phase by incubation with 12.5 mM hydroxyurea (HU) for 4 h at 28°C, followed by a release into fresh HU-free medium. Denatured total cell extracts (TCA extracts) were prepared at 60 and 90 min post-release, resolved via 5.5% SDS-PAGE, and analyzed by Western blot using an anti-GFP antibody. Total protein visualization by Coomassie blue staining was used as a loading control.

**Figure S6.**
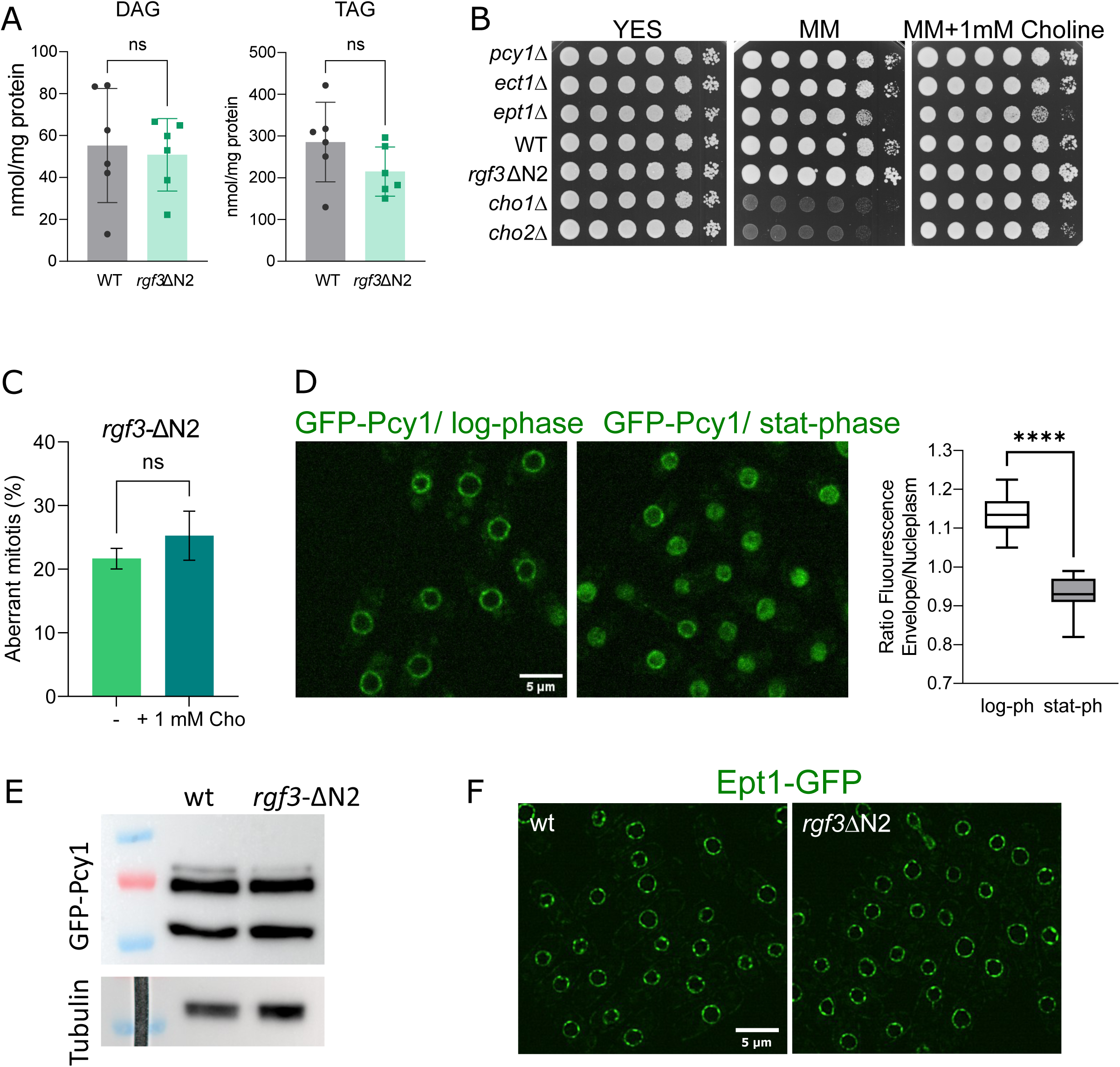
Lipidomic quantification of neutral lipids, choline-dependent genetic rescues, and expression profiles of Kennedy pathway enzymes. **(A)** Lipidomic analysis of wild-type (WT) and *rgf3*ΔN2 cells showing the relative abundance (nmol/mg protein) of diacylglycerol (DAG) (left) and triacylglycerol (TAG) (right). Data represent the mean ± SD from n = 6 biological replicates. Statistical analysis was performed using a two-tailed Student’s *t*-test assuming equal variance (n.s., non-significant). **_(B)_** Choline-dependent growth rescue assays in CDP–DAG and Kennedy pathway mutant backgrounds. Cells from the indicated strains (*pcy1*Δ, *ect1*Δ, *ept1*Δ, WT, *rgf3*ΔN2, *cho1*Δ, and *cho2*Δ) were cultured in YES medium, adjusted to an initial O.D._600_ of 2.7, and a series of three two-fold serial dilutions followed by two ten-fold serial dilutions were spotted onto YES, minimal medium (MM), and MM supplemented with 1 mM choline solid plates. Plates were imaged after 2–3 days of incubation at 28°C. **(C)** Quantitative analysis of the percentage of aberrant mitosis in log-phase *rgf3*ΔN2 cells cultured in the presence (+) or absence (–) of 1 mM choline added 3 h prior to time-lapse imaging. Data represent the mean ± SD from three independent experimental replicates (n > 20 cells analyzed per condition). Statistical analysis was judged by a Student’s *t*-test (n.s., non-significant). **(D)** Morphometric quantification of the nuclear envelope-to-nucleoplasm GFP-Pcy1 fluorescence ratio. WT cells expressing GFP-Pcy1 were analyzed during exponential growth (O.D._600 ∼_0.6) or stationary phase (O.D._600 ∼_9) (n = 39 cells analyzed per condition). The box plot defines the interquartile range (IQR); whiskers represent Tukey’s range, and the horizontal line inside the box indicates the median value. Statistical significance was determined using a Student’s *t*-test (****, p< 0.0001). **(E)** Western blot analysis of GFP-Pcy1 protein expression levels. Denatured total cell extracts (TCA extracts) were prepared from exponentially growing WT and *rgf3*ΔN2 cells expressing GFP-Pcy1. Samples were resolved via SDS-PAGE and immunoblotted using an anti-GFP antibody. Detection of α-tubulin was utilized as an internal loading control. **(F)** Representative fluorescence micrographs displaying the subcellular localization of Ept1-GFP in exponentially growing WT and *rgf3*ΔN2 mutant strains. Scale bar, 5 µm.

